# Multidimensional MRI reveals cellular-scale microstructural phenotypes in human brain aging

**DOI:** 10.64898/2026.03.29.715117

**Authors:** Joon Sik Park, Eppu Manninen, Shunxing Bao, Bennett A. Landman, Yihong Yang, Dan Benjamini

**Author notes:** **Corresponding author:** Dan Benjamini, Ph.D., Multiscale Imaging and Integrative Biophysics Unit, National Institute on Aging, NIH, Baltimore, Maryland, USA.

## Abstract

Brain aging is accompanied by profound cellular and microstructural changes that precede overt tissue loss, yet in vivo MRI studies largely emphasize macroscopic measures or isolated diffusion and relaxation metrics, providing limited insight into how cellular-scale tissue architecture is altered across the adult lifespan. Here, we apply multidimensional diffusion–relaxation MRI (MD-MRI) to map voxel-wise microstructural phenotypes in cognitively unimpaired adults spanning early adulthood to late life (23–77 years). Rather than relying on predefined compartment models, MD-MRI resolves continuous voxel-wise distributions in a joint diffusion–relaxation space, enabling an integrated, model-free description of how cellular shape, size, restriction, and chemical environment vary with age. Using this approach, we reveal age-related multilateral shifts within a complex microstructural landscape, marked by increasing heterogeneity and disorder across brain tissue. With age, cellular-scale features showed a systematic transition from small to larger length-scale structures accompanied by reduced microscopic restriction, indicating a loss of fine cellular barriers and expansion of extracellular space. In parallel, we show tissue-dependent alterations in fast-relaxing properties aligned with known gray- and white-matter aging processes, including iron accumulation and myelin loss. Together, these findings indicate that normative brain aging involves progressive reorganization of structure and composition at the cellular level, rather than uniform shifts in bulk tissue properties. By decoupling cellular-scale shape, size, restriction, and chemical environment in vivo, MD-MRI identifies increasing cellular heterogeneity and breakdown of microscopic restriction as central features of human brain aging and provides a biologically interpretable framework for linking microstructural reorganization to age-related functional decline.

## 1. Introduction

Age-related alterations in brain structure occur across multiple spatial scales and influence cognitive and functional trajectories throughout adulthood (Raz and Rodrigue, 2006). At the macroscopic level, structural MRI studies consistently demonstrate cortical thinning and progressive volume loss in both superficial and deep gray matter (GM), with many of these changes accelerating during midlife and continuing into later decades (Resnick et al., 2003; Storsve et al., 2014). White matter (WM) undergoes parallel degeneration, including reduced tract orientational coherence, decreased myelination (Qian et al., 2020), and disruptions across projection, association, and commissural pathways (Schilling et al., 2022). Together, these findings highlight widespread structural remodeling of the aging brain, but largely reflect late-stage or cumulative changes.

At the microscopic level, postmortem histological studies reveal a richer and more complex picture of normative brain aging. Hallmark features include dendritic regression, synaptic pruning, glial remodeling, and gradual expansion of the extracellular space, reflecting coordinated yet heterogeneous alterations in cytoarchitecture and tissue microstructure (Peters, 2002; Dickstein et al., 2013; Popov et al., 2021). These cellular-level processes are thought to emerge early and to contribute to age-related changes in neural function, long before overt tissue loss becomes apparent.

Diffusion MRI (dMRI) provides one of the primary noninvasive tools for interrogating brain microstructure by measuring water motion constrained by cellular and extracellular boundaries (Callaghan, 2011). Conventional models such as diffusion tensor imaging (DTI) (Basser et al., 1994) robustly demonstrate age-related increases in mean diffusivity and decreases in fractional anisotropy (Schilling et al., 2022). More recent biophysical models, including NODDI (Zhang et al., 2012) and SANDI (Palombo et al., 2020), aim to increase biological specificity and suggest age-related reductions in neurite density (Lee et al., 2024), often interpreted as reflecting demyelination, neurite loss, or extracellular space expansion. However, these models rely on strong a priori assumptions about tissue composition and geometry, limiting their validity in heterogeneous or weakly anisotropic regions such as cortical gray matter (Jelescu et al., 2020; Afzali et al., 2021). As brain aging involves multiple concurrent and spatially overlapping microstructural processes, collapsing this complexity into a small set of summary parameters may obscure subtle or early changes critical for understanding normative aging.

Growing evidence suggests that microstructural degeneration precedes measurable reductions in cortical or subcortical volume, even in cognitively unimpaired individuals (Rathi et al., 2014; Singh et al., 2024). This temporal dissociation underscores the need for noninvasive imaging approaches capable of detecting subtle and spatially heterogeneous microstructural changes before overt anatomical decline occurs. Approaches that resolve intra-voxel heterogeneity are particularly valuable, as brain tissue contains diverse water environments that vary in cellular density, membrane properties, extracellular content, diffusion behavior, and relaxation characteristics (Syková and Nicholson, 2008; Qian et al., 2020). Such heterogeneity cannot be adequately captured by unidimensional imaging measures but is essential for characterizing latent tissue remodeling during normal aging.

Recent technological and theoretical advancements have led to an imaging framework that can address these limitations by merging two important aspects: simultaneous sensitivity to multiple MR contrasts and the ability to resolve their distribution within tissue. This strategy, known as multidimensional MRI (MD-MRI), departs from conventional diffusion or relaxation imaging, which assigns each voxel a single averaged value, by acquiring data linked to multiple properties—typically diffusion tensors together with longitudinal (R_1_) and transverse (R_2_) relaxation rates—and reconstructing how these parameters are distributed rather than collapsing them into a mean value (Benjamini, 2020; Slator et al., 2021a). In practice, MD-MRI can separate contributions from water pools with distinct microstructural phenotypes without enforcing specific compartment models, reducing partial-volume effects and isolating environments that scalar maps blend together (English et al., 1991). Hardware and sequence innovations have made these multidimensional experiments feasible in vivo and ex vivo (Benjamini and Basser, 2016; Kim et al., 2017; de Almeida Martins and Topgaard, 2018; Kundu et al., 2023), enabling applications that range from characterizing microstructural variation in healthy neural tissue (Benjamini and Basser, 2017; de Almeida Martins et al., 2021) to detecting changes associated with inflammation, degeneration, and tumor biology (Naranjo et al., 2021; Benjamini et al., 2020; Slator et al., 2023). Notably, ex vivo MD-MRI has uncovered glial-related alterations in Alzheimer’s disease that are not detectable with conventional diffusion or relaxation imaging (Barsoum et al., 2025).

Building on the MD-MRI framework that combines R_1_ and R_2_ relaxometry with tensor-valued diffusion encoding (Martin et al., 2021), a newly developed and efficient in vivo MD-MRI protocol has recently been introduced (Johnson et al., 2024; Manninen et al., 2024). This approach enables measurement of the frequency-dependent diffusion spectrum, **D**(*ω*), which probes time-dependent diffusion restrictions shaped by cellular morphology (Narvaez et al., 2022). In this study, we leverage this novel MRI framework to chart the microstructural landscape of the aging brain across the adult lifespan. Using data from 46 cognitively unimpaired adults aged 23–77 years, we derive microstructural empirical “phenotype” based on cellular-level shape (microscopic anisotropy), size (isotropic diffusivity), and chemical environment (R_2_ relaxation), along with restriction (diffusion frequency dependence). Because these phenotypes do not depend on biophysical modeling, they can be quantified across any brain region, as demonstrated here in cortical GM, deep GM nuclei, and major WM pathways. Our central hypothesis is that aging drives a systematic and diverse redistribution of diffusion-relaxation subdomains. We find age-related changes across multiple tissue characteristics: shifts from small toward both intermediate and large length-scale phenotypes in GM, high-to-medium anisotropy phenotypes shift in WM, and low-to-high R_2_ relaxation phenotypes shift in both cortex and subcortex. Together, this framework enables a unified, cellular-scale description of brain aging by framing age-related changes as a coordinated redistribution of microstructural features, thereby linking in vivo MRI measurements to cellular processes known to underlie normative brain aging.

## 2. Methods

### 2.1. Participants

A total of 46 cognitively unimpaired healthy participants (20-77 years; mean age ± SD = 47.3 ± 17.6 years; 23 females) were drawn from ongoing control cohorts at the National Institute on Drug Abuse (NIDA). Participants were stratified into three age groups for descriptive and visualization purposes: young (20-31 years), middle-aged (32-57 years), and aged adults (61-77 years), as detailed in Table 1. All individuals provided informed consent prior to participation in the study, which was approved by the NIDA Institutional Review Board. Exclusion criteria included major medical illness or current medication use, a history of neurological or psychiatric disorders, or any history of substance abuse.

**Table 1.**
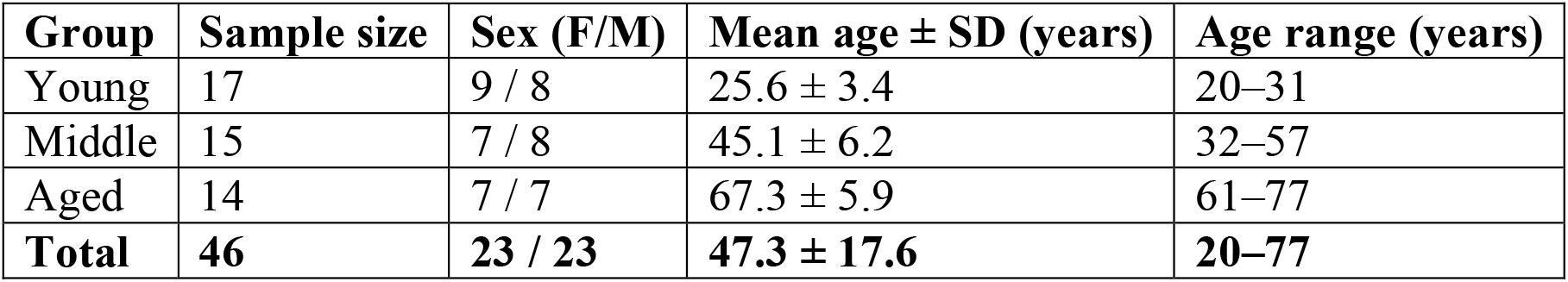
Demographic characteristics of the participants.

| **Group** | **Sample size** | **Sex (F/M)** | **Mean age ± SD (years)** | **Age range (years)** |
| --- | --- | --- | --- | --- |
| Young | 17 | 9 / 8 | 25.6 ± 3.4 | 20–31 |
| Middle | 15 | 7 / 8 | 45.1 ± 6.2 | 32–57 |
| Aged | 14 | 7 / 7 | 67.3 ± 5.9 | 61–77 |
| **Total** | **46** | **23 / 23** | **47.3 ± 17.6** | **20–77** |

### 2.2 Data Acquisition

MRI data were acquired on a 3T MAGNETOM Prisma scanner (Siemens Healthineers, Erlangen, Germany) using a 32-channel receive array. Multidimensional diffusion-relaxometry measurements were obtained with a single-shot spin-echo EPI sequence incorporating tensor-valued diffusion encoding generated by numerically optimized free gradient waveforms. Images were acquired at 2 mm isotropic resolution (FOV = 228×228×110 mm^3^, bandwidth=1512 Hz/pixel), with GRAPPA acceleration (factor=2, 24 reference lines), an effective echo spacing of 0.8 ms, partial Fourier 6/8, and axial slice orientation (Johnson et al., 2024).

The diffusion-encoding scheme followed a sparse yet information-rich sampling strategy. In addition to a non-diffusion-weighted (b = 0) volume, the protocol included linear (*b*_Δ_= 1), spherical (*b*_Δ_ = 0), and planar (*b*_Δ_ = −0.5) b-tensors spanning b-values from 0.1 to 3 ms/μm^2^. Each waveform was characterized by a well-defined diffusion spectral profile, with centroid frequencies (*ω*_*cent*_/2π) ranging from 6.6 to 21 Hz, enabling sensitivity to time-dependent diffusion across biologically relevant frequency bands (Yon et al., 2025).

Relaxation sensitivity was achieved by varying repetition times (TR = 0.62, 1.75, 3.5, 5, 7, and 7.6 s) and echo times (TE = 40, 63, 83, and 150 ms), resulting in 139 unique diffusion-relaxation encoding combinations. The total MD-MRI acquisition time was approximately 40 minutes. Images were acquired with an anterior-to-posterior phase-encoding direction, and an additional reversed (*b* = 0) posterior-to-anterior volume was collected to enable susceptibility-induced distortion correction. Further details are provided in the Supplementary Information and in Fig. S1.

High-resolution structural reference images were acquired using a fat-suppressed *T*_1_-weighted MPRAGE sequence (TR = 1900 ms, TE = 3.42 ms, 1 mm isotropic resolution). These anatomical images were used for co-registration, brain extraction, and region-of-interest definition.

### 2.3. Data Preprocessing

MD-MRI data were preprocessed following current recommendations for sparse MD-MRI imaging (Johnson et al., 2024). All diffusion-weighted volumes were first concatenated and denoised using the Marchenko-Pastur principal component analysis (MP-PCA) method (Veraart et al., 2016). Motion and eddy-current distortions were corrected using the DIFFPREP module of the TORTOISE software package (Pierpaoli et al., 2010), employing a physically based quadratic transformation model and a normalized mutual information similarity metric. Susceptibility-induced geometric distortions were corrected with the DR-BUDDI algorithm (Irfanoglu et al., 2015) by providing the acquired anterior-posterior (AP) and posterior-anterior (PA) phase-encoding b = 0 images together with a synthetic *T*_2_-weighted (b = 0-like) image generated from the *T*_1_-weighted anatomical scan (Schilling et al., 2019).

### 2.4 Multidimensional Data Processing

Preprocessed MD-MRI data were analyzed in MATLAB (MathWorks, Natick, MA) using a Monte Carlo inversion framework implemented in the multidimensional diffusion MRI toolbox (Martin et al., 2021; Narvaez et al., 2022). The measured signal was modeled as a weighted superposition of discrete components spanning the joint diffusion-relaxation parameter space:

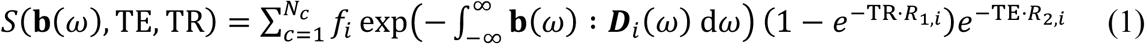

where *f*_*i*_ denotes the weight of the i-th component and : represents the generalized scalar product. Each frequency-dependent diffusion tensor ***D***_*i*_(*ω*) was modeled as an axisymmetric Lorentzian function parameterized by axial diffusivity *D*_∥,*i*_, radial diffusivity *D*_⊥, *i*_, high-frequency isotropic diffusivity *D*_0,*i*_, orientation angles (*θ*_*i*_, *ϕ*_*i*_), axial and radial transition frequencies (Γ_∥,*i*_ Γ_⊥,*i*_), and longitudinal and transverse relaxation rates (*R*_1,*i*_, *R*_2,*i*_) (de Almeida Martins and Topgaard, 2018; Narvaez et al., 2024; Johnson et al., 2024). For convenience in interpretation, axial and radial diffusivities are expressed here as isotropic diffusivity, *D*_*iso*_, = (*D*_∥_ + 2*D*⊥) / 3, and squared microscopic diffusion anisotropy, 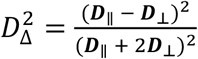. We computed voxel-wise means and variances for all diffusion-relaxation parameters, denoted as E[·] and V[·] (e.g., E[*D*_*iso*_]), respectively. Following conventions, diffusion frequency dependence was quantified by finite-difference slopes computed across the sampled centroid frequency range (i.e., between *ω*_*min*_ = 6.6 Hz and *ω*_*max*_ = 21 Hz), noted as Δ_*ω*/2π_ E[*D*_*iso*_] and 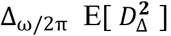for isotropic diffusivity and anisotropy, respectively (Narvaez et al., 2024; Johnson et al., 2024). Full technical details regarding implementation are provided in the Supplementary Information.

### 2.5 Definition of diffusion-relaxation subdomains

By treating **D*(*ω*)***-*R*_1_,-*R*_2_ distributions as spectra, images of individual spectral components can be obtained through integration over predefined diffusion–relaxation parameter subdomains. The resulting value, ranging from 0 to 1, represents the signal fraction of that component and can be computed voxel-wise to produce spatial maps of specific microstructural “phenotypes”, based on their length-scale, shape, and chemical microenvironment. This approach has previously been used to probe microscopic eccentricity (Benjamini and Basser, 2014), diffusivity–*T*_2_ subdomains (Benjamini and Basser, 2017; Pas et al., 2020; Slator et al., 2021b; Benjamini et al., 2022) and diffusivity–anisotropy subdomains in vivo (de Almeida Martins et al., 2020; Martin et al., 2021; Manninen et al., 2024). Here, we combine these strategies by partitioning the continuous dimensions of 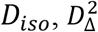 and *R*_2_ into subdomains, based on heuristic principles:

- The isotropic diffusivity in MD-MRI is commonly partitioned into two subdomains (de Almeida Martins et al., 2020). Here, we expand that division into three subdomains: low (0.05 < *D*_*iso*_< 1.5 μm^2^/ms), intermediate (1.5 < *D*_*iso*_ < 3 μm^2^/ms), and high (*D*_*iso*_ > 3 μm^2^/ms) spectral subdomains, which capture small, intermediate, and large microscopic length-scales, respectively.
- The diffusion microscopic anisotropy in MD-MRI is commonly partitioned into two subdomains (de Almeida Martins et al., 2020). Here, we expand that division into three subdomains: low 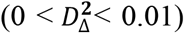, intermediate 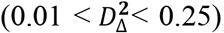, and high 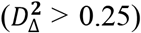 spectral subdomains, which capture low, intermediate, and high microscopic anisotropy, respectively.
- The *R*_2_ relaxation is normally not partitioned in MD-MRI. Here, we partition it to include low (1 < *R*_2_ < 20 s^−1^) and high (*R*_2_ > 20 s^−1^) spectral subdomains, which capture contributions from tissue water and macromolecule-bound water, respectively (Alonso-Ortiz et al., 2015).

In total, we define and investigate 10 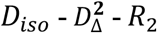 subdomains, which we term here “microstructural phenotypes”. For convenience, we use the left superscripts *𝓁, 𝓂*, and *𝓀* to denote low, intermediate, and high ranges along each parameter dimension, respectively. For example, the phenotype 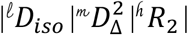 represents the normalized signal fraction of a microstructural phenotype with low isotropic diffusivity, intermediate diffusion anisotropy, and high *R*_2_ relaxation. For the ^𝓀^*D*_*iso*_ range, we include the full anisotropy range as previously established (de Almeida Martins et al., 2020; Martin et al., 2021; Manninen et al., 2024), and denote it as 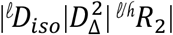. Further details are provided in the Supplementary Information.

These microstructural phenotypes are diffusion-frequency dependent. However, throughout most of this work we analyze them at the lowest diffusion frequency (*ω*_*min*_ = 6.6 Hz) and therefore omit the *ω*_*min*_ notation for clarity. When explicitly examining frequency dependence, we use *ω*_*min*_ and *ω*_*max*_ to indicate the frequencies at which the phenotypes were derived.

### 2.6 Image registration and whole brain segmentation

Cortical and subcortical GM regions were obtained using the spatially localized atlas network tiles (SLANT) method (Huo et al., 2019), which applies region-specific 3D convolutional networks following standard preprocessing. The resulting segmentations yield 132 anatomical labels based on the BrainCOLOR atlas and have demonstrated high reproducibility. GM segmentation was performed on 1 mm *T*_1_-weighted images in native space and downsampled to 2 mm resolution to match the MD-MRI data.

White matter tract labels were derived from the Johns Hopkins University (JHU) DTI-based WM atlas using Advanced Normalization Tools (ANTs v2.5.2). Because direct registration from individual participant images to atlas space was inaccurate, age-specific group templates were used as an intermediate step. Participants were assigned to one of three age groups (Table 1), and for each group a template was generated by co-registering *T*_1_ -weighted images using affine and symmetric normalization (SyN). Templates were down-sampled to 2 mm isotropic resolution to match the MD-MRI images and the JHU atlas fractional anisotropy map. Group-averaged 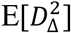 images were registered to the JHU atlas using affine and SyN transforms. The resulting transforms—from participant to group template and from group template to atlas space—were concatenated, inverted, and applied to map atlas labels into each participant’s native space, with a single interpolation used to minimize interpolation errors.

To enable voxel-wise comparisons, MD-MRI parameter maps were transformed to FSL’s MNI152 template space. Group-registered *T*_1_ -weighted images (1 mm isotropic) were first registered to the MNI152 *T*_1_-weighted template (1 mm isotropic) using affine and SyN transforms. The resulting transforms—from participant space to group template and from group template to MNI152 space—were concatenated and applied to the MD-MRI maps, with a single interpolation used to minimize interpolation errors.

In this study we chose to focus on a wide array of cortical regions of interest (ROI) that included the frontal pole (FRP), anterior orbital gyrus (AOrG), posterior orbital gyrus (POrG), opercular part of the inferior frontal gyrus (OpIFG), triangular part of the inferior frontal gyrus (TrIFG), middle frontal gyrus (MFG), medial precentral gyrus (MPrG), supplementary motor cortex (SMC), anterior insula (AI), superior temporal gyrus (STG), postcentral gyrus (PoG), posterior cingulate gyrus (PCgG), precuneus (PCu), angular gyrus (AnG), middle occipital gyrus (MOG), cuneus, and entorhinal cortex (EntC). Subcortical ROIs included the hippocampus, caudate, putamen, and thalamus. White matter tracts included the genu of the corpus callosum (genu CC), anterior corona radiata (ACR), superior corona radiata (SCR), sagittal stratum (SS), external capsule (EC), cingulum–hippocampus bundle (CH), and the superior fronto-occipital fasciculus (SFOF). Left and right ROIs were combined to create single regions. In total, 28 ROIs were investigated.

### 2.7 Statistical Analysis

Statistical analyses were performed in MATLAB (MathWorks, Natick, MA) to assess age-related effects on MD-MRI microstructural phenotypes across cortical, subcortical, and WM ROIs. Age associations were evaluated using a quadratic regression model applied independently to each MD-MRI parameter and ROI. Sex was included as a covariate using effect coding (male = 0.5, female = −0.5). The regression model was defined as 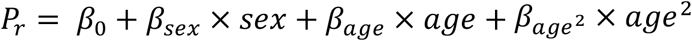, where *P*_*r*_ denotes the ROI-averaged MD-MRI parameter of interest (e.g., 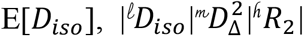, etc.) for region r. Linear age effects were quantified by *β*_*age*_, while nonlinear lifespan trajectories were assessed via the quadratic term 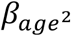. All continuous variables were z-standardized prior to regression, such that regression coefficients represent standardized effect sizes.

Multiple-comparison correction across all ROIs and MD-MRI metrics was performed using the Benjamini–Hochberg false discovery rate (FDR) procedure, with statistical significance defined as *p*FDR < 0.05 (Storey, 2002).

## 3. RESULTS

### 3.1 Interpretation framework for microstructural phenotypes

Unlike biophysical compartment models such as NODDI or SANDI—which link each parameter to predefined geometries (e.g., spheres for soma, sticks for neurites) and a fixed number of intra-voxel compartments—MD-MRI estimates the distribution of diffusion-relaxation components directly from the measured signal, without assuming any specific tissue geometry or compartment count. Consequently, the derived parameters should be interpreted as phenomenological descriptors of diffusion and relaxation behavior, rather than as direct quantitative measures of specific cellular constituents.

While MD-MRI provides voxel-wise joint distributions of frequency-dependent diffusion tensors **D**(*ω*) and relaxation parameters *R*_1_, and *R*_2_,, we choose here to focus on a projection onto the 3D parameter space spanned by the isotropic diffusivity, *D*_*iso*_, the squared microscopic diffusion anisotropy, 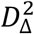, and the transverse relaxation rate *R*_2_,. Together, these three parameters define a multidimensional diffusion-relaxation space in which distinct signal components occupy characteristic subdomains. Hence, we define the term “microstructural phenotype” to describe the normalized signal fraction from predefined ranges within the 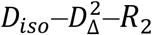, distribution. As detailed in the Methods Section, we partitioned the continuous dimensions of *D*_*iso*_, 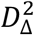, and *R*_2_, into three, three, and two subdomains, based on heuristic principles. Here, the left superscripts 𝓁, 𝓂, *and* 𝓀 denote low, intermediate, and high ranges along each parameter dimension, respectively, as illustrated in Fig. 1A. For example, the phenotype 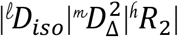 represents the normalized signal fraction of a microstructural phenotype with low isotropic diffusivity, intermediate diffusion anisotropy, and high *R*_2_, relaxation.

**Figure 1.**
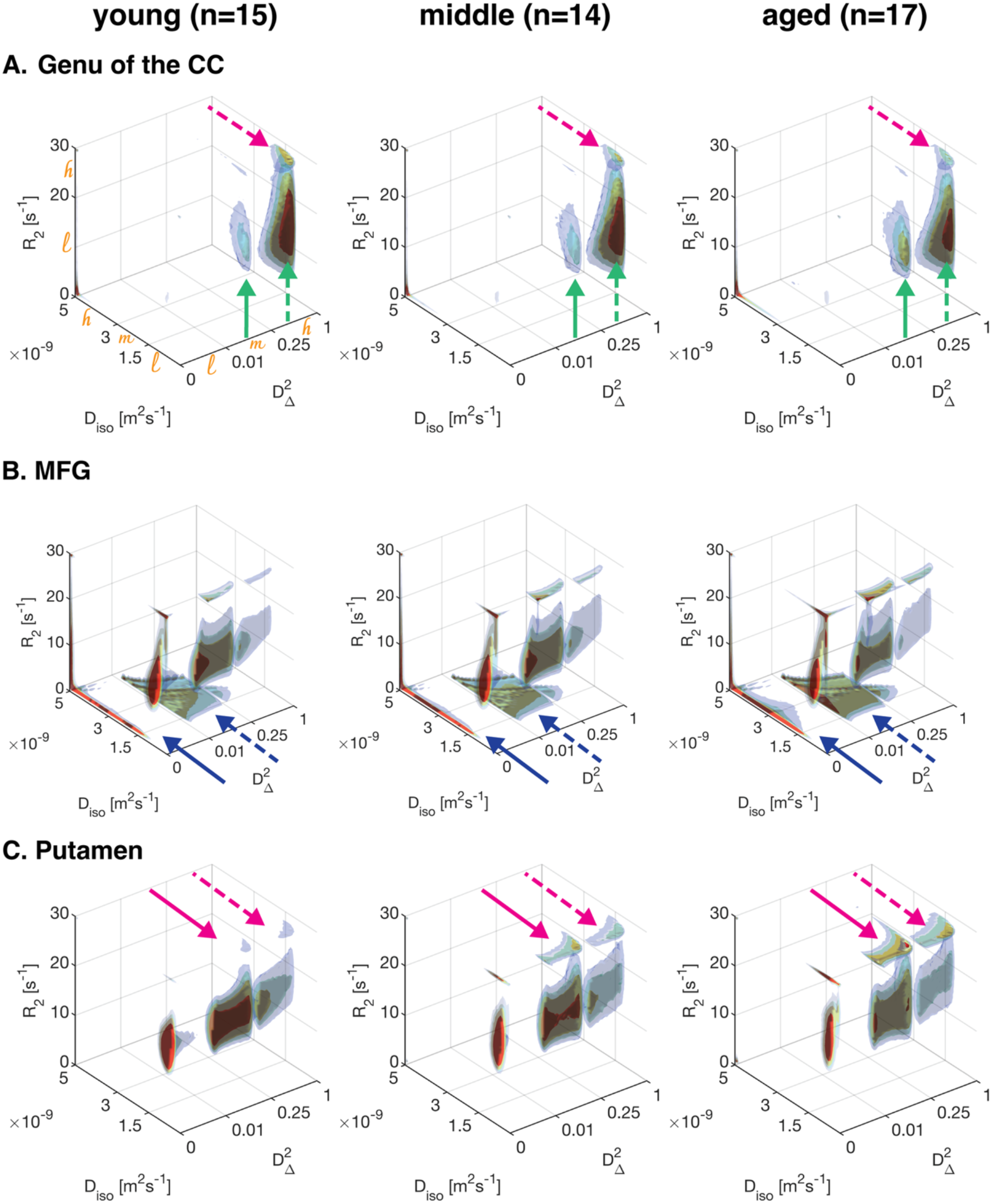
Age-related redistribution of joint diffusion-relaxation phenotypes across major brain tissue classes. Group-averaged three-dimensional joint 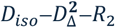, distributions projected at minimal frequency, *ω*_min_= 6.6 Hz, are shown for young (left), middle-aged (middle), and aged adults (right) in the (A) genu of the corpus callosum, (B) middle frontal gyrus (MFG), and (C) putamen. Across brain regions, aging is marked by multilateral microstructural shifts that reflect complex trajectories and together form a shared aging signature. Low, intermediate, and high phenotype ranges, l, m, and h, are indicated in (A). Arrows indicate major age-related shifts in microstructural phenotypes.

Importantly, because these microstructural phenotypes sum to 1 within each voxel, any age-related effects can be assessed and interpreted as systematic redistributions across phenotypes within the joint 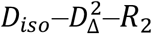, space, rather than as changes in predefined biological compartments.

### 3.2 Multilateral diffusion–relaxation shifts reveal complex age-related microstructural change

The participants were grouped into three categories reflecting different phases of adult life: young adults (n = 15), middle-aged adults (n = 14), and aged adults (n = 17). Figure 1 illustrates group differences and age effects in microscopic length scale, shape, and chemical composition reflected in the joint 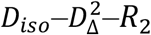, distributions projected on the lowest frequency *ω*_*min*_ = 6.6 Hz, for representative brain regions spanning major tissue classes, including a commissural WM tract (genu of the corpus callosum), cortical GM (middle frontal gyrus), and subcortical GM (putamen), shown in Fig. 1A, B, and C, respectively. These regions were selected to illustrate how age-related microstructural alterations manifest across tissues with distinct organizational architectures, ranging from highly anisotropic fiber bundles to heterogeneous cortical and subcortical regions.

In the genu of the corpus callosum (Fig. 1A), younger adults exhibited a dominant diffusion-relaxation phenotype characterized by low isotropic diffusivity, high microscopic diffusion anisotropy (dashed green arrow), and a fast *R*_2_ component (dashed pink arrow), occupying a compact region of the joint space. With increasing age, this distribution systematically shifts toward the intermediate anisotropy 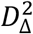 range (solid green arrow), and reduced intensity of the fast *R*_2_, component (dashed pink arrow), indicating a progressive reduction in microstructural restriction and directional coherence. Notably, this change was expressed as a shift of the dominant phenotype 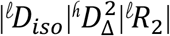 towards 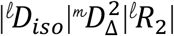, rather than the emergence of a qualitatively new diffusion-relaxation component.

The middle frontal gyrus region (Fig. 1B) showed broader, more heterogeneous distributions across all age groups, consistent with the inherently less ordered cortical microarchitecture. We observed a complex age-related pattern across microstructural phenotypes: small length-scale phenotypes (^𝓁^*D*_*iso*_) across all shapes shifted toward intermediate and large diffusivity ranges, mainly within low and intermediate anisotropy ranges (solid and dashed blue arrows, respectively). In addition, a buildup of the small length-scale and fast *R*_2_ phenotypes can be seen.

In the putamen (Fig. 1C), redistribution followed a distinct pattern, with prominent age-related increases in the small length-scale and fast *R*_2_ phenotypes (dashed and solid pink arrows for high and intermediate anisotropy, respectively). A concomitant age-related decrease in low *R*_2_ phenotypes was observed, along with a general shift from higher to lower microstructural anisotropy. This shift was more pronounced along the relaxation dimension than in cortical GM, suggesting that age-related changes in the macromolecular environment and water–tissue interactions may play a particularly important role in age-related microstructural remodeling within deep GM nuclei.

Across all three tissue classes, aging was characterized by coordinated redistributions within the joint 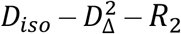 space rather than isolated changes in individual parameters. These multilateral shifts reflect complex microstructural trajectories with age, constituting a shared aging signature.

The corresponding distributions evaluated at the highest diffusion frequency (*ω*_*max*_ = 21 Hz) are shown in the Supplementary Information (Fig. S2), illustrating how microscopic length scale, shape, and chemical composition vary with diffusion frequency across the same representative white- and gray-matter regions. An explicit frequency dependence analysis is provided in Section 3.7.

### 3.3 Age group differences in diffusion-relaxation parameter maps

Multidimensional MRI delivers voxel-wise distributions like the ones seen in Fig. 1. The information can be most straightforwardly summarized by mapping weighted averages and variances across the different dimensions. Such group-averaged maps are shown in Fig. 2A, along with corresponding aged-to-young percentage difference maps (Fig. 2B), where E[·] and V[·] denote the voxel-wise average and variance of a given dimension, respectively. In addition, Δ_*ω*/2π_ E[·] captures the diffusion frequency dependence of the corresponding diffusion metrics.

**Figure 2.**
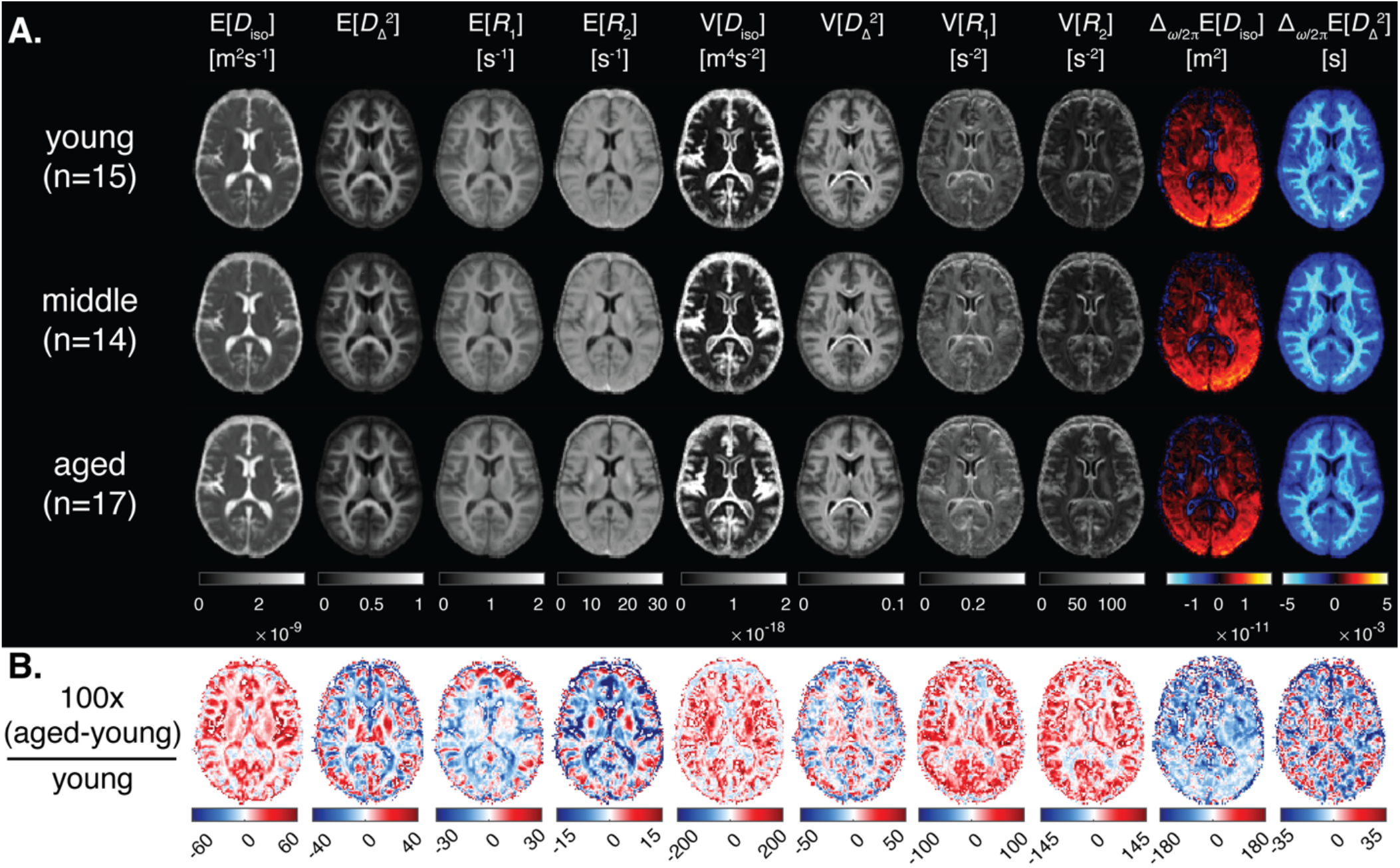
Diffusion–relaxation parameters shown in (A) as participant-averaged maps for each age group. Panel (B) presents percent change maps for each metric, highlighting differences between the young (20–32 years) and aged (61–77 years) groups. Displayed parameters include expectation values E[D_iso_], 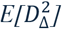, E[R_1_] and E[R_2_], corresponding to MD, microscopic anisotropy, 1/T_2_, and 1/T_2_; the corresponding variances V[D_iso_], 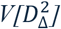, V[R_1_], and V[R_2_], which capture intra-voxel parameter heterogeneity; and the diffusion-frequency slopes Δ_*ω*/2π_E[D_iso_] and 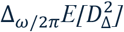, which reflect microstructural restriction.

The average parameter maps E[*D*_*iso*_], 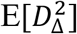, E[*R*_1_] and E[*R*_2_] roughly correspond to DTI’s mean diffusivity, microscopic diffusion anisotropy, 1/*T*_1_, and 1/*T*_2_. Visual inspection shows that these metrics follow the expected age-related trajectories, in which older adults exhibit higher mean isotropic diffusivity and lower microscopic anisotropy. Mean relaxometry parameters showed opposing age-related trends in GM and WM, increasing with age in GM and decreasing in WM (Fig. 2B).

Beyond differences in mean values, variance maps (i.e., V[·]) demonstrated pronounced age-related increases in all parameters, particularly in frontal and parietal association cortices, reflecting heightened intra-voxel heterogeneity and greater dispersion of microstructural environments in aging tissue. Complementing these findings, diffusion frequency slopes, Δ_*ω*/2π_E[*D*_*iso*_], and 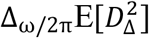, exhibited systematic flattening with age, suggestive of attenuated restriction-driven frequency dependence and increased extracellular contributions.

Region-of-interest–averaged MD-MRI diffusion–relaxation parameters, stratified by age group, are given in Fig. S3.

### 3.4 Age group differences in microstructural phenotype maps

The most distinctive and powerful feature of MD-MRI is its ability to resolve diffusion– relaxation subdomains within a voxel, yielding effective intra-voxel microstructural resolution. Maps of these subdomains directly visualize how signal weight redistributes across distinct regions of the joint diffusion–relaxation space. Rather than representing changes in predefined model-based biological compartments, they capture age-related shifts in the relative contributions of different diffusion–relaxation subdomains. Importantly, these quantities sum to 1 as they represent the normalized signal fraction from all subdomains. These microstructural phenotypes are shown in Fig. 3, which reveals pronounced age-related redistribution of signal contributions within the joint diffusion–relaxation space. Because these novel microstructural phenotypes follow opposing age-related trajectories in cortical and subcortical GM and WM regions, we next examine each of them separately.

**Figure 3.**
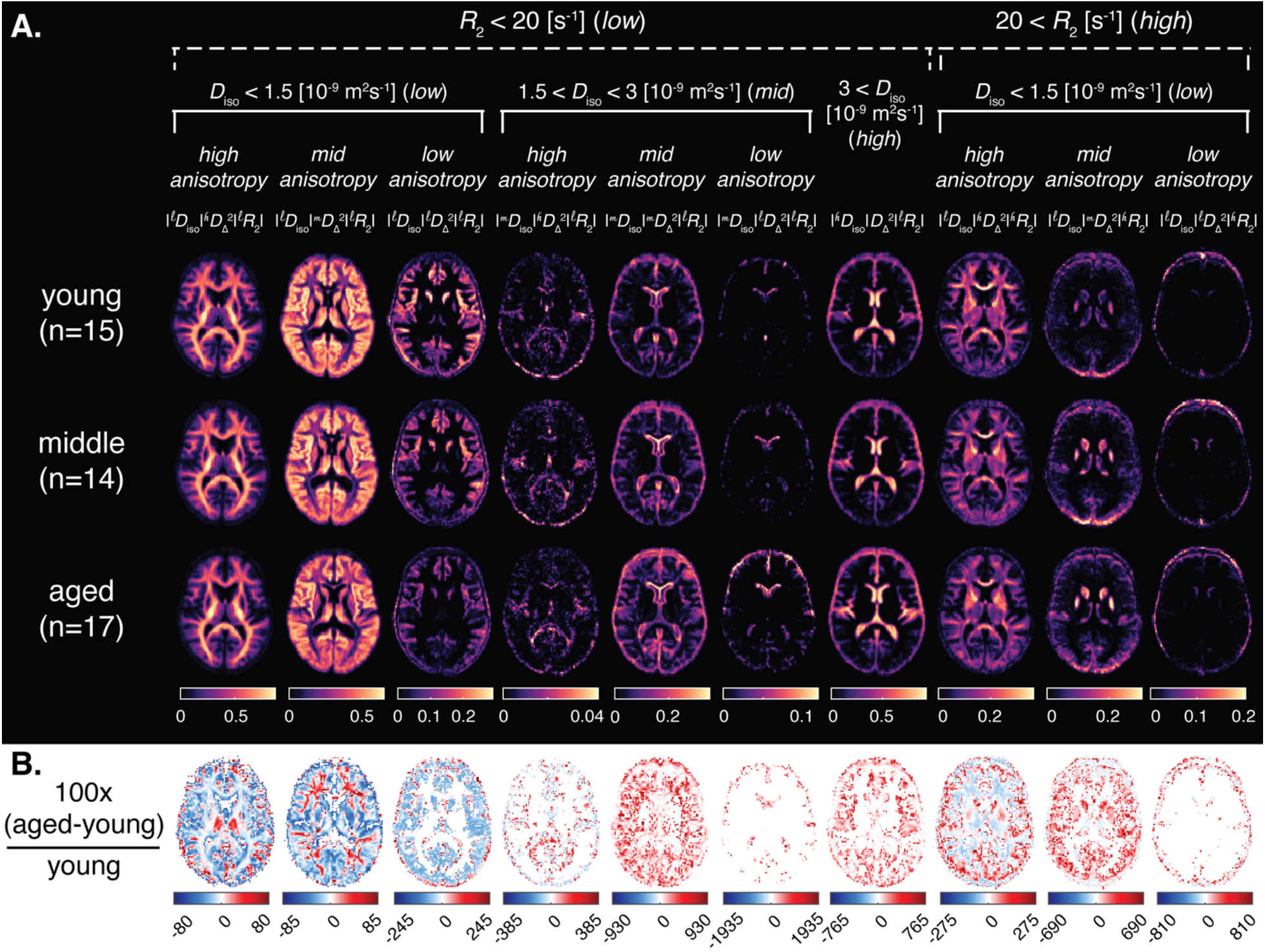
Microstructural phenotypes shown in (A) as participant-averaged maps for each age group. Panel (B) presents percent change maps for each metric, highlighting differences between the young (20–32 years) and aged (61–77 years) groups. These phenotypes represent the normalized signal contributions of distinct subdomains of the joint 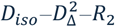, space, defined by combinations of low (𝓁), intermediate (𝓂), and high (𝓀) values of isotropic diffusivity, diffusion anisotropy, and R_2_ relaxation rate, without assuming correspondence to specific biological compartments. Group-difference maps in (B) highlight widespread age-related redistribution toward larger cellular length-scales, increased macromolecular and iron content in GM, and reduced microscopic anisotropy.

In cortical and subcortical GM regions, aging was accompanied by an increase in the characteristic cellular length-scale, whereas microscopic anisotropy was preserved. This pattern is reflected by negative aged-to-young relative differences in 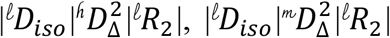, and 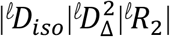, alongside concomitant positive aged-to-young relative differences in 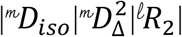, and 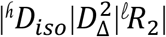 (Fig. 3B). Similarly, tissue phenotypes with fast *R*_2_ relaxation and small cellular length-scales show an increasing prevalence with age across the GM, indicative of increased macromolecular and iron content. The caudate and putamen show especially strong effects, with percent differences as high as 690%.

In WM, microstructural redistribution is more constrained, characterized by a decrease in highly anisotropic, small length-scale tissue phenotypes and concurrent increases in intermediate-anisotropy, small length-scale and unrestricted, high-diffusivity phenotypes. Specifically, 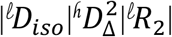 shifts toward 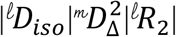, and 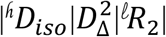. In addition, the prevalence of small length-scale, high-anisotropy, high-*R*_2_ phenotypes is reduced, consistent with decreased macromolecular content (e.g., myelin).

Region-of-interest–averaged MD-MRI microstructural phenotypes, stratified by age group, are given in Fig. S4. Some phenotypes have clear regional specificity; for example, both 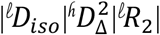 and 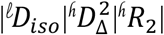 are more prevalent in WM. Conversely, lower anisotropy subdomains, e.g., 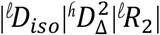 or 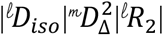, mainly dominate cortical and subcortical regions (Fig. S4).

### 3.5 Linear and nonlinear age associations in diffusion-relaxation metrics

To quantify these observations, we first examined both linear and quadratic associations with age (centered at 47.6 years) across several diffusion–relaxation MD-MRI parameters. Associations were expressed using z-scored (standardized) coefficients, *β*_*age*_ and 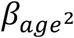, which represent the change in the MR variable, in units of standard deviation, per year. The resulting age-related effect sizes are shown in Fig. 4A and B for the linear and quadratic associations, respectively. In addition, scatter plots from representative regions and MD-MRI parameters are shown in Fig. 4C.

**Figure 4.**
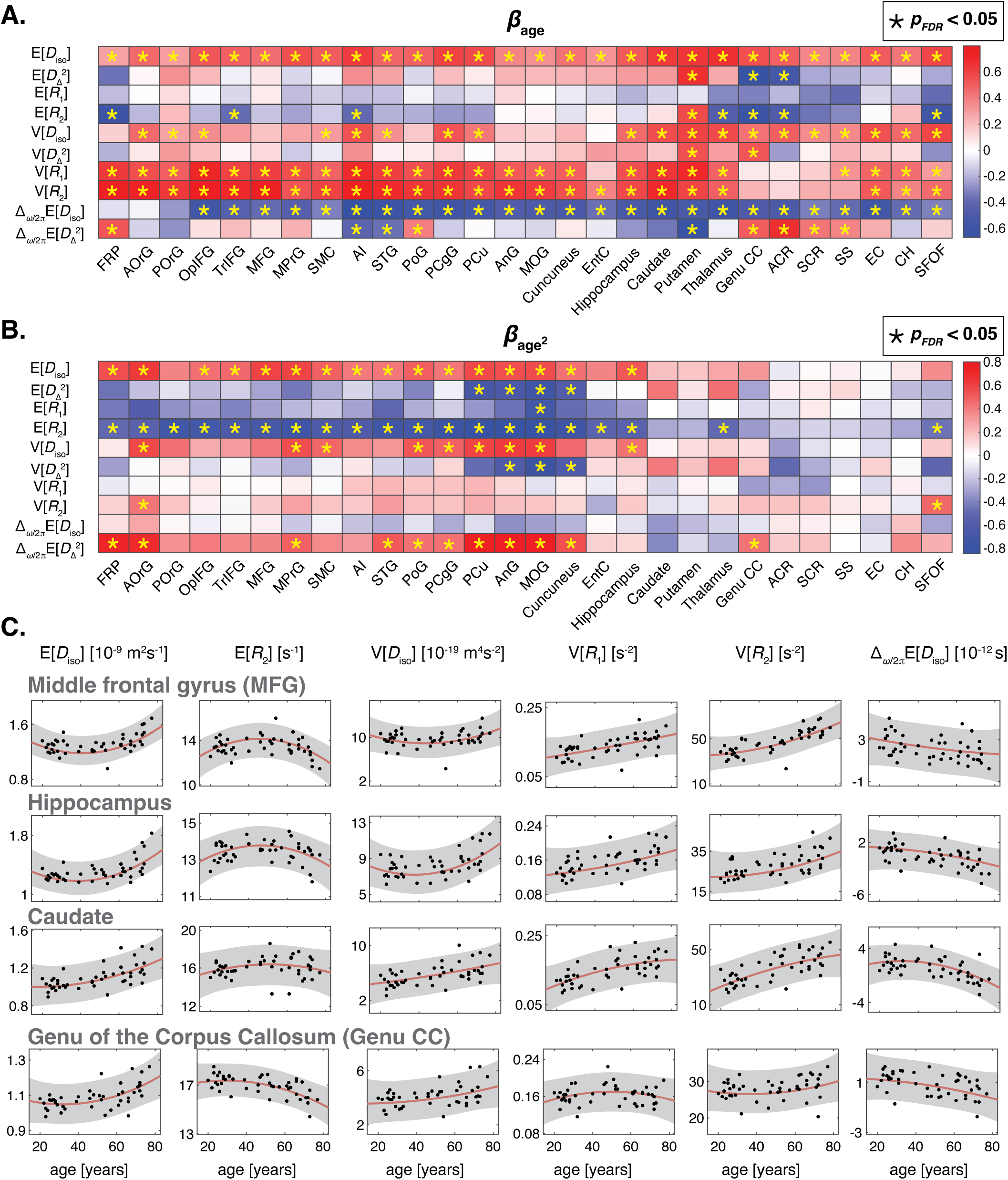
Linear and nonlinear age associations of multidimensional MD-MRI parameters. (A) Standardized linear age coefficients (β_age_) for expectation values, variances, and diffusion-frequency slopes of diffusion–relaxation parameters across cortical, deep GM, and WM ROIs. (B) Corresponding quadratic age coefficients 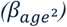 capturing nonlinear lifespan effects. Asterisks indicate false-discovery-rate corrected significance (p_FDR_ < 0.05). Scatter plots from representative regions and MD-MRI parameters are shown in panel (C). Solid lines denote model fits, with shaded regions indicating 95% confidence intervals.

Consistent with their distinct microstructural organization, GM and WM regions exhibited divergent age-related trends in diffusion–relaxation MD-MRI parameters. All regions showed positive linear age associations with isotropic diffusivity, E[*D*_*iso*_]; however, significant quadratic associations, after correction for multiple comparisons, were restricted to GM regions (Fig. 4). Diffusion anisotropy, 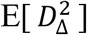, showed positive linear age associations in GM and negative associations in WM, although most did not survive FDR correction. Transverse relaxation, E[*R*_2_], exhibited weak negative linear age associations across most regions, while demonstrating strong inverted U-shaped quadratic relationships in GM.

In addition to changes in average parameter values, variance measures exhibited strong and spatially extensive age effects. Variances of isotropic diffusivity and relaxation rates, particularly V[*D*_*iso*_] and V[*R*_2_], increased linearly with age across association cortices, limbic regions, deep nuclei, and major WM tracts (Fig. 4). These findings indicate increasing intra-voxel heterogeneity and a broader dispersion of diffusion–relaxation environments in older adults, extending beyond shifts in average tissue properties.

Strong negative linear associations with age were observed for the diffusion frequency dependence of isotropic diffusivity, Δ_*ω*/2π_ E[*D*_*iso*_], across all regions (Fig. 4). These findings suggest a progressive reduction in restriction-driven frequency dependence with age, consistent with decreased microstructural complexity and greater extracellular water contributions.

### 3.6 Linear and nonlinear age associations in microstructural phenotypes

To quantify age-related redistribution of diffusion–relaxation properties, we examined region-wise linear and nonlinear associations between age and microstructural phenotypes across cortical, deep GM, and WM ROIs (Fig. 5). For each region, regression models evaluated linear (*β*_*age*_) and quadratic 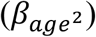 age effects while controlling for sex, with significance assessed using false-discovery-rate correction. The resulting z-scored coefficients indicate per-year changes in phenotype in standard deviation units. Because these microstructural phenotypes represent normalized signal fractions that sum to one, they are inherently interconnected, and their age trajectories are best interpreted together.

**Figure 5.**
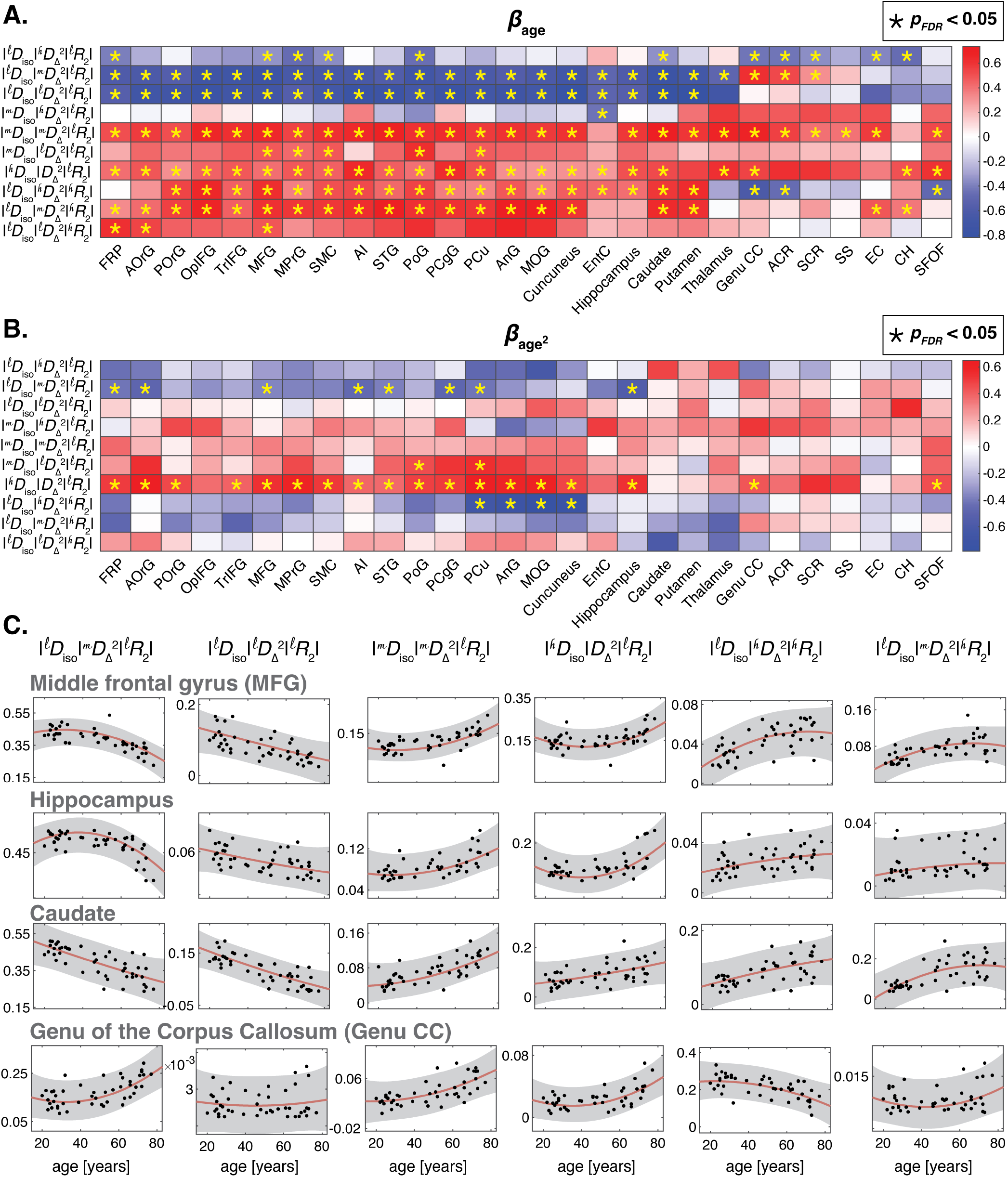
Linear and nonlinear age associations of microstructural phenotypes across brain regions. (A) Standardized linear age coefficients (β_age_) estimated for signal fractions associated with distinct diffusion–relaxation subdomains across cortical, deep GM, and WM ROIs. (B) Corresponding quadratic age coefficients 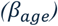 capturing nonlinear lifespan trajectories. Scatter plots from representative regions and MD-MRI parameters are shown in panel (C). Solid lines denote model fits, with shaded regions indicating 95% confidence intervals. Phenotypes with ROI-averaged intensity under 0.01 were omitted from the plot to avoid bias.

When examining the low *R*_2_ subdomains, GM regions generally exhibited age-related decreases in phenotypes characterized by small cellular length-scales (i.e., ^𝓁^ *D*_*iso*_), with corresponding increases in phenotypes associated with intermediate and large length-scales (i.e., ^𝓂^*D*_*iso*_ and ^𝓁^*D*_*iso*_). Only two phenotypes, 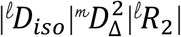 and 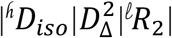, exhibited strong and consistent quadratic age relationships (Fig. 5A and B). Microstructurally, these observations are consistent with decreased presence of cellular barriers and greater extracellular water contributions with age.

In WM, redistribution is limited in low *R*_2_ subdomains, with negative linear associations in highly anisotropic, small length-scale phenotypes and positive associations in intermediate-anisotropy and unrestricted, high-diffusivity phenotypes. Accordingly, 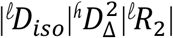 shifts toward 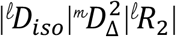 and 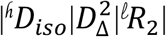. From a microstructural perspective, these findings reflect reduced cellular barriers, increased orientational dispersion, and greater extracellular water contributions with aging.

Finally, we examined the high *R*_2_ subdomains, which are sensitive to macromolecular and iron content. Several age-related patterns emerged. First, all high *R*_2_ phenotypes were characterized by small cellular length scales. Second, significant positive linear age associations were confined to high- and intermediate-anisotropy phenotypes and were observed primarily in GM, consistent with iron accumulation. Third, negative linear age associations were detected in the genu of the corpus callosum, anterior corona radiata, and superior fronto-occipital fasciculus, consistent with myelin loss.

### 3.7 Diffusion frequency dependence of microstructural phenotypes

Up to this point, all reported microstructural phenotypes were evaluated at the lowest diffusion frequency. Time- or frequency-dependent diffusion arises from microscopic restrictions. As such, it provides important biophysical context and interpretability, motivating the investigation of frequency-dependent effects in this work. In the following analysis, we explicitly examine the dependence of microstructural phenotypes on diffusion frequency, reporting differences between the highest (*ω*_*max*_=21 Hz) and lowest (*ω*_*min*_=6.6 Hz) frequencies.

Figure 6 summarizes the diffusion frequency dependence of the microstructural phenotypes. Panels A and B show the absolute and relative differences, respectively, between high- and low-frequency estimates averaged over the young adult group. Differences are separated and color-coded by brain region (cortical, subcortical, and WM).

**Figure 6.**
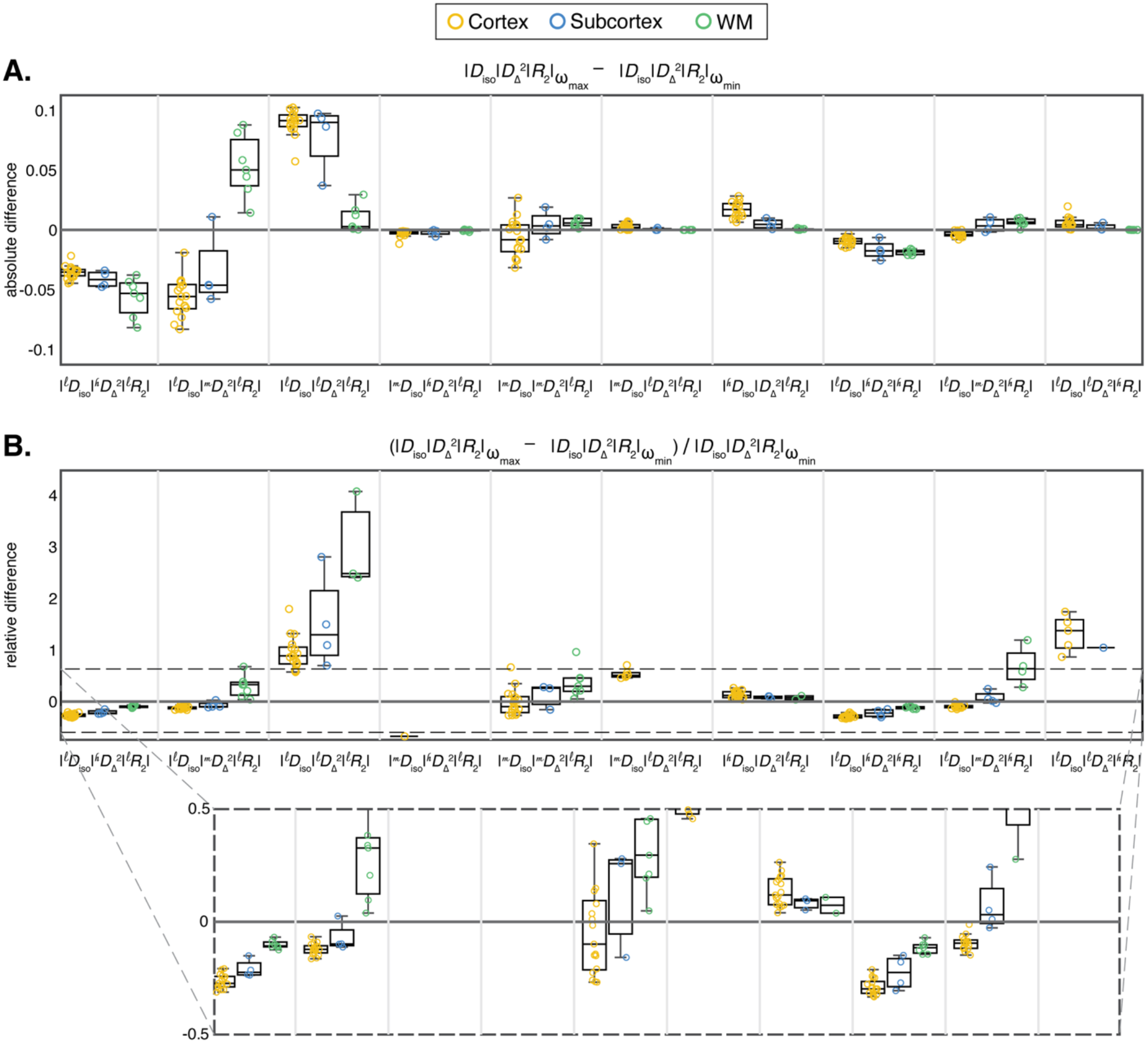
Diffusion frequency dependence of microstructural phenotypes. Panels A and B show the absolute and relative differences, respectively, between high- and low-frequency estimates averaged across the young adult group. Differences are grouped and color-coded by brain region (cortical GM, subcortical GM, and WM). Phenotypes with ROI-averaged intensity under 0.01 were omitted from the plot to avoid bias.

Considerable frequency dependence was observed in several microstructural phenotypes. The biggest effect was observed in the near-isotropic, small length-scale parameters, namely, 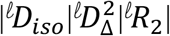 and 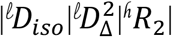. In both cases, higher diffusion frequencies are associated with larger signal fractions (up to 300% difference in WM, Fig. 6B). The high anisotropy, small length-scale parameters, 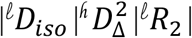 and 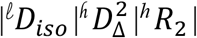, also exhibited strong restriction-driven frequency dependence. Meanwhile, the large length-scale parameter, 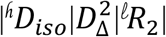, showed very little relative change with frequency, pointing to a lack of restriction.

The corresponding analysis for the aged adult group is provided in the Supplementary Information (Fig. S5), illustrating absolute and relative differences between high- and low-frequency estimates and confirming the presence of diffusion frequency–dependent effects. The age associations of the relative differences parameter are shown in Fig. S6.

## 4. Discussion

Using MD-MRI, this cross-sectional study mapped novel microstructural phenotypes in a cohort of cognitively unimpaired adults spanning a broad age range and revealed age-related multilateral shifts within a complex microstructural landscape. Rather than focusing on isolated diffusion or relaxation metrics, the MD-MRI framework resolves continuous voxel-wise distributions in a joint diffusion–relaxation space, enabling an integrated description of how cellular shape, size, restriction, and chemical composition reorganize with age. Our results indicate that cellular-scale shape and restriction play an important role in brain aging, with small length-scale structures exhibiting near-isotropic or intermediate anisotropy remodeling into larger length-scale structures. In parallel, fast-relaxing R_2_ phenotypes exhibiting cellular-scale restriction increase with age in GM and decrease in WM, likely reflecting iron accumulation and myelin loss, respectively.

Popular biophysical compartment models such as SANDI and NODDI assign parameters to predefined cellular structures by assuming a simplified microstructural model *a priori* and then fitting the data to estimate the relative contributions of each compartment. In contrast, MD-MRI inverts this paradigm by first reconstructing continuous diffusion–relaxation distributions directly from the measured signal, without imposing assumptions about compartment number or tissue geometry, and subsequently deriving empirical “phenotypes” that reflect distinct microstructural cellular features.

Prior MRI studies reported microstructural simplification in the aging brain, including increased diffusivity, reduced anisotropy, and altered relaxation properties across WM regions (Schilling et al., 2022; Qian et al., 2020). Recent GM-focused diffusion studies similarly found decreased cellular density and increased microstructural heterogeneity with age (Lee et al., 2024; Singh et al., 2025; Bouhrara et al., 2023; Singh et al., 2024). These results can be interpreted as different projections of a common underlying redistribution of diffusion–relaxation parameters. Extending this perspective, we show here that aging is marked by coordinated shifts within a joint diffusion–relaxation parameter space (Fig. 1), rather than by the appearance of distinct new signal components. The diffusion–relaxation phenotypes identified here therefore provide a compact and interpretable summary of dominant signal states, without implying a one-to-one correspondence with specific cellular or histological compartments.

The age-related patterns observed in conventional diffusion and relaxation parameters (Figs. 2 and 4) are consistent with prior reports. In particular, the strong quadratic U-shaped association between age and E[*D*_*iso*_], comparable to the DTI-derived mean diffusivity, has been previously described (Bouhrara et al., 2023; Singh et al., 2024). Similarly, an inverted U-shaped age dependence of E[*R*_2_] has also been reported (Hagiwara et al., 2021). While conventional diffusion and relaxation MRI parameters have been extensively investigated in the context of aging, MD-MRI uniquely enables the characterization of intra-voxel variance. Beyond changes in mean parameter values, the observed age-related increases in variance across diffusion and relaxation metrics suggest a progressive rise in intra-voxel microstructural heterogeneity with aging, which could be driven by changes in extracellular volume and the presence of reactive astrocytes and microglia. The prominence of these effects in frontal and parietal association cortices points to an increasing dispersion of cellular and tissue environments in regions known to be particularly vulnerable to age-related reorganization (Resnick et al., 2003).

Another distinctive feature of this study is the ability to characterize the frequency dependence of E[*D*_*iso*_] and 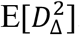. Because diffusion frequency dependence is directly linked to microscopic restriction (Yon et al., 2025), the observed age-related reductions in Δ_*ω*/2π_E[*D*_*iso*_] together with predominantly positive linear and quadratic age associations in 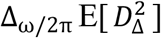 across cortical, subcortical, and WM regions point to a systematic alteration of cellular-scale barriers with aging. Collectively, these patterns are consistent with a progressive breakdown of microstructural restriction, reflecting age-related weakening or reorganization of cellular boundaries rather than uniform changes in bulk diffusivity.

A central strength of MD-MRI is its ability to resolve diffusion–relaxation subdomains within a voxel, providing effective intra-voxel microstructural information. We defined in this study 10 subdomains in the joint 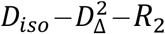 space, which capture different aspects of cellular-level morphology and chemical environment. The resulting maps, shown in Fig. 3, quantify the normalized signal fraction from each of the subdomains and, importantly, sum to 1. Age-related redistributions within low-*R*_2_ diffusion–relaxation subdomains point to systematic reorganization of cellular-scale microstructure with aging (Fig. 5). In GM, the shift away from phenotypes associated with small cellular length scales toward intermediate and larger length-scale components can be hypothesized to reflect a reduction in effective cellular barriers and an increased contribution of extracellular water. The presence of quadratic age relationships in select phenotypes further suggests non-monotonic remodeling processes rather than simple linear degradation. In WM, redistribution within low-*R*_2_ subdomains was more constrained but followed a coherent pattern, with decreases in highly anisotropic, small–length-scale phenotypes accompanied by increases in intermediate-anisotropy and less restricted components. This pattern is indicative of increased orientational dispersion and extracellular expansion, consistent with age-related loosening of microstructural organization rather than uniform loss of fiber architecture.

Distinct age-related effects emerged within high-*R*_2_ subdomains, which are sensitive to macromolecular density and iron content. The restriction of high-*R*_2_ phenotypes to small cellular length scales across tissues suggests that these signal components predominantly reflect compact microstructural environments. Positive age associations observed primarily in anisotropic GM phenotypes are consistent with progressive iron accumulation (Burgetova et al., 2021), whereas negative age associations localized to anterior WM tracts, including the genu of the corpus callosum and corona radiata, align with known patterns of age-related myelin loss (Qian et al., 2020). Together, these findings indicate that high-*R*_2_ subdomains capture tissue-specific aging processes that are distinct from the extracellular and barrier-related changes observed in low-*R*_2_ phenotypes.

The strong diffusion frequency dependence observed across specific diffusion–relaxation phenotypes provides insight into age-related changes in microstructural restriction imposed by cellular membranes and barriers. The pronounced frequency sensitivity of near-isotropic, small length-scale phenotypes (Fig. 6) suggests that these signal components arise from water pools confined by dense arrangements of cellular membranes and other restricting structures, whose contribution becomes more prominent when diffusion probes shorter length- and time-scales. The magnitude of these effects, particularly in WM, highlights the value of frequency-dependent measurements for capturing age-related alterations in the prevalence and organization of restricting barriers. Similarly, the frequency dependence observed in highly anisotropic, small–length-scale phenotypes is consistent with restriction within elongated, membrane-bounded structures such as axons or glial processes. In contrast, the relative frequency insensitivity of large length-scale, high-diffusivity phenotypes supports their interpretation as reflecting weakly restricted or extracellular environments, which are expected to contribute more prominently as restricting cellular barriers become less dominant with aging. Together, these findings indicate that diffusion frequency encoding provides a sensitive means of probing age-related reorganization of membrane-associated microstructural barriers.

The coordinated redistribution of diffusion–relaxation signal states observed here is consistent with multiple, partially overlapping biological processes known to accompany normative brain aging. Histological and ultrastructural studies have documented age-related dendritic regression (Dickstein et al., 2013), synaptic pruning (Morrison and Baxter, 2012), and glial remodeling across cortical and subcortical regions (Popov et al., 2021), together with gradual expansion of the extracellular space, all of which are expected to reduce microscopic restriction and directional coherence. Within this context, age-related shifts in WM—including changes in microscopic diffusion anisotropy, attenuation of diffusion-frequency dependence, and alterations in high-*R*_2_ phenotypes—can be hypothesized to reflect reorganization of axonal packing, myelin-associated structure, and glial contributions. In cortical GM, the broadening of diffusion–relaxation distributions and the expansion of intermediate- and high-Diso subdomains can be interpreted as increasing heterogeneity of the neuropil, where dendritic simplification, synapse loss, and astrocytic remodeling may coexist within individual voxels. Deep GM regions exhibited a distinct aging signature marked by increased high-*R*_2_ phenotype prevalence and reduced microscopic diffusion anisotropy; given the complex architecture of these regions, including the presence of crossing fibers, these changes are consistent with reduced neurite orientation dispersion and a decline in crossing fiber contributions, with potential implications for regional connectivity.

Several limitations should be considered when interpreting these findings. First, the cross-sectional design, spanning early adulthood to late life, limits inference about within-subject aging trajectories and cannot fully separate true aging effects from cohort-related influences; longitudinal MD-MRI studies will be necessary to establish the temporal progression and rate of microstructural redistribution. Second, although the MD-MRI framework enables model-free reconstruction of continuous diffusion–relaxation distributions, the resulting phenotypes represent phenomenological signal states rather than direct proxies for specific cellular or histological compartments. While the observed patterns are consistent with known biological processes, definitive mechanistic attribution will require integration with complementary imaging modalities and, ultimately, postmortem validation. Third, the MD-MRI protocol reflects a trade-off between multidimensional sensitivity and clinical feasibility, with an acquisition time of approximately 40 minutes; in particular, the narrow frequency window of 6.6–21 Hz presents a limiting factor for observing restriction. Further optimization of frequency encoding range (Yon et al., 2025) and inversion strategies (Park et al., 2025) may improve sensitivity and robustness. Finally, the present analyses were restricted to cognitively unimpaired adults, and extending this framework to pathological aging and neurodegenerative populations will be essential to determine the specificity and clinical relevance of diffusion–relaxation phenotypes.

## 5. Conclusion

Using MD-MRI, we demonstrate that normative brain aging is characterized by coordinated redistributions within a joint diffusion–relaxation parameter space, rather than by the emergence of discrete new microstructural features. This multidimensional framework reveals age-related increases in intra-voxel heterogeneity alongside phenotype-specific shifts, including a redistribution from small to larger effective length scales, reduced signatures of microscopic restriction, and tissue-dependent changes in fast-relaxing signal components. Together, these patterns are consistent with progressive reorganization of cellular-scale barriers, increased extracellular contributions, and distinct gray- and white-matter aging processes, including iron accumulation and myelin loss. By resolving diffusion–relaxation features and incorporating diffusion-frequency dependence, MD-MRI provides an integrated, model-free description of how cellular-scale size, shape, restriction, and chemical environment reorganize with age, complementing conventional diffusion and relaxometry measures. Although longitudinal and multimodal validation will be required to establish biological specificity, these findings position multidimensional diffusion–relaxation phenotyping as a promising approach for linking cognitive observations to cellular-scale mechanisms of brain aging.

## AUTHOR CONTRIBUTIONS

Study design: DB, YY. Study conduct and data collection: JSP, YY, DB. Data analysis: JSP, EM, SB, BAL, DB. Data interpretation: JSP, DB. Drafting the manuscript: JSP, DB.

## ACKNOWLEDGMENTS

This research was supported by the Intramural Research Program of the National Institutes of Health (NIH). The contributions of the NIH author(s) are considered Works of the United States Government. The findings and conclusions presented in this paper are those of the author(s) and do not necessarily reflect the views of the NIH or the U.S. Department of Health and Human Services.

## DATA AVAILABILITY STATEMENT

Code to process the MD-MRI data is freely available as implemented in the multi-dimensional diffusion MRI toolbox (https://github.com/markus-nilsson/md-dmri). The data used in the study are available upon direct request as well as the conditions for its sharing or re-use.

