## Supplementary for "Multidimensional MRI reveals cellular-scale microstructural phenotypes in human brain aging"

### Supplementary Methods

#### MD-MRI data acquisition

**MRI system and pulse sequence:** All MRI data were acquired on a 3T MAGNETOM Prisma system (Siemens Healthineers, Erlangen, Germany) using a 32-channel receive-only head coil. Multidimensional diffusion–relaxometry MRI (MD-MRI) data were collected using a single-shot spin-echo echo-planar imaging (EPI) sequence modified for tensor-valued diffusion encoding (TDE) with numerically optimized free gradient waveforms [1, 2].

**Acquisition parameters:** MD-MRI was acquired at 2-mm isotropic spatial resolution with the following parameters: field of view (FOV) = $228 \times 228 \times 110$ mm$^3$, voxel size = $2 \times 2 \times 2$ mm$^3$, acquisition bandwidth = 1512 Hz/pixel, in-plane parallel imaging using GRAPPA with acceleration factor $R = 2$ (24 reference lines), effective echo spacing = 0.8 ms, partial Fourier factor = 0.75 in the phase-encoding direction, and axial slice orientation. A comprehensive summary of acquisition parameters is provided in supplementary Table 1.

**Protocol design:** The MD-MRI protocol comprised 139 unique diffusion–relaxation measurements with an overall scan time of approximately 40 minutes per participant. The protocol was derived from a previously reported 633-point $(D(\omega), R_1, R_2)$ acquisition [1],with encoding parameters empirically selected to reduce protocol dimensionality while preserving sensitivity to diffusion-relaxation contrasts and extending the ranges of repetition time (TR) and echo time (TE) relative to the original design.

**Diffusion-relaxation encoding:** Diffusion encoding was achieved using numerically optimized gradient waveforms [3] designed to generate linear ($b_\Delta = 1$), spherical ($b_\Delta = 0$), and planar ($b_\Delta = -0.5$) b-tensors, with b-values ranging from 0.1 to 3 ms/$\mu$m$^2$, in addition to one non-diffusion-weighted ($b = 0$ ms/$\mu$m$^2$) volume. Sensitivity to longitudinal and transverse relaxation was achieved by sampling multiple repetition times and echo times, with TR = (0.62, 1.75, 3.5, 5, 7, 7.6) s and TE = (40, 63, 83, 150) ms.

The diffusion gradient waveforms used in the acquisition and their corresponding diffusion spectra are shown in Supplementary Fig. S1. The applied waveforms exhibited broad spectral content characterized by centroid frequencies $\omega_{\text{cent}}/2\pi$ spanning 6.6–21 Hz. Each waveform contains a distribution of frequencies, and the centroid frequency is reported only as a representative descriptor. All subsequent signal modeling and inversion steps explicitly used the full spectral representation $b(\omega)$ rather than a single-frequency approximation. The maximum frequencies reached depended on the b-value, with approximate maxima of 21, 15, and 11 Hz for $b$ = 0.5, 1.5, and 3 ms/$\mu$m$^2$, respectively.

To enable correction of susceptibility-induced geometric distortions, MD-MRI data were acquired with a single anterior-to-posterior (AP) phase-encoding direction together with an additional reversed posterior-to-anterior (PA) $b = 0$ volume.

**Structural imaging:** In addition to MD-MRI, fat-suppressed $T_1$-weighted MPRAGE images were acquired for anatomical reference and ROI definition (TR = 1900 ms, TE = 3.42 ms, 1-mm isotropic resolution). Structural images were used exclusively for image registration and anatomical segmentation.

#### MD-MRI data processing

**Data preprocessing:** MD-MRI data were preprocessed following recent recommendations for sparse multidimensional diffusion–relaxometry MRI acquisitions [4]. All diffusion-weighted volumes were first concatenated and denoised using the

| Parameter | Value / Description |
|---|---|
| Scanner | 3T MAGNETOM Prisma (Siemens Healthcare, Erlangen, Germany) |
| Head coil | 32-channel receive-only |
| Structural scan | Fat-suppressed $T_1$-weighted MPRAGE (1 mm isotropic) |
| Sequence | Single-shot spin-echo EPI with tensor-valued diffusion encoding |
| Voxel size | 2 mm isotropic |
| Field of view (FOV) | $228 \times 228 \times 110$ mm$^3$ |
| Bandwidth | 1512 Hz/Px |
| Parallel imaging | GRAPPA (R = 2), 24 reference lines |
| Echo spacing | 0.8 ms |
| Partial Fourier | 0.75 (phase-encoding direction) |
| Phase encoding | AP (plus one reversed PA $b = 0$ for distortion correction) |
| Repetition times (TR) | 0.62, 1.75, 3.5, 5, 7, 7.6 s |
| Echo times (TE) | 40, 63, 83, 150 ms |
| Diffusion weighting ($b$) | 0.1–3 ms/$\mu$m$^2$ |
| B-tensor shapes | Linear ($b_\Delta = 1$), Planar ($b_\Delta = -0.5$), Spherical ($b_\Delta = 0$) |
| Frequency range | Centroid $\omega_{\text{cent}}/2\pi = 6.6$–21 Hz |
| Number of volumes | 139 unique diffusion-relaxation measurements |
| Scan time | 40 minutes |

Supplementary Table 1: Comprehensive summary of the in vivo MD-MRI acquisition protocol.

Marchenko–Pastur principal component analysis (MP-PCA) method [5]. For partial Fourier acquisitions, Gibbs ringing correction was applied using a local subvoxel-shift approach [6, 7].

Motion- and eddy-current-induced distortions were corrected using the DIFFPREP module of the TORTOISE software package [8], employing a physically based parsimonious quadratic transformation model together with a normalized mutual information similarity metric. Susceptibility-induced geometric distortions were corrected using the DR-BUDDI framework [9]. For this purpose, the acquired anterior–posterior (AP) and posterior–anterior (PA) phase-encoding $b = 0$ images were combined with a synthetic $T_2$-weighted image generated from the $T_1$-weighted anatomical scan [10].

All preprocessing steps were combined into a single final interpolation step, yielding fully corrected MD-MRI volumes aligned to anatomical space at native MD-MRI resolution (2-mm isotropic). No visible slice-to-volume motion or motion-induced signal dropouts were observed in this dataset; therefore, these correction options were not applied.

**Signal modeling and Monte Carlo inversion:** Preprocessed MD-MRI data were analyzed in MATLAB (MathWorks, Natick, MA) using a Monte Carlo (MC) inversion framework implemented in the multidimensional diffusion MRI toolbox [11, 12, 13, 14]. The MD-MRI signal was modeled as a weighted superposition of discrete components spanning the joint diffusion-relaxation parameter space according to

$$S(b(\omega), \text{TE}, \text{TR}) = \sum_{i=1}^{N_c} f_i \exp\left(-\int_{-\infty}^{\infty} b(\omega) : \mathbf{D}_i(\omega)\, d\omega\right)\left(1 - e^{-\text{TR}\cdot R_{1,i}}\right) e^{-\text{TE}\cdot R_{2,i}}, \quad (1)$$

where $f_i$ denotes the signal fraction of the $i$-th component and $N_c$ is the number of fitted components.

Each frequency-dependent diffusion tensor $\mathbf{D}_i(\omega)$ was approximated as an axisymmetric Lorentzian parameterized by axial and radial diffusivities $(D_{\|,i}, D_{\perp,i})$, orientation angles $(\theta_i, \phi_i)$, high-frequency isotropic diffusivity $D_{0,i}$, axial and radial transition frequencies $(\Gamma_{\|,i}, \Gamma_{\perp,i})$, and longitudinal and transverse relaxation rates $(R_{1,i}, R_{2,i})$ [15, 16, 17, 4]. These parameters were sampled within biologically plausible bounds: $0.05 \le D_\|, D_\perp, D_0 \le 5\ \mu\text{m}^2/\text{ms}$; $0 \le \theta \le \pi$, $0 \le \phi \le 2\pi$; $0.2 \le R_1 \le 2\ \text{s}^{-1}$, $1 \le R_2 \le 30\ \text{s}^{-1}$; and $0.01 \le \Gamma_\|, \Gamma_\perp \le 10^4\ \text{s}^{-1}$. Axial and radial diffusivities were reparameterized as isotropic diffusivity

$$D_{\text{iso}} = \frac{D_\| + 2D_\perp}{3}, \quad (2)$$

and squared microscopic diffusion anisotropy

$$D_\Delta^2 = \frac{(D_\| - D_\perp)^2}{(D_\| + 2D_\perp)^2}. \quad (3)$$

The MC inversion was performed using nonnegative least squares combined with quasi-genetic filtering and bootstrapping with replacement to address the ill-conditioned nature of the inverse problem [17, 13]. For each voxel, $N_b = 100$ bootstrap realizations were generated, each producing up to $N_c = 10$ discrete components. Algorithmic parameters were set to $N_{\text{in}} = 200$, $N_p = 20$, and $N_m = 1$. As a result, for each imaging voxel, a total of weighted signal components were estimated across bootstrap realizations, providing a robust empirical sampling of the underlying diffusion-relaxation signal. The resulting voxel-wise solutions were evaluated at selected values of $\omega/2\pi$ within the experimentally sampled frequency range (6.6-21 Hz), yielding frequency-dependent representations of the diffusion-relaxation signal in the $(D_{\mathrm{iso}}(\omega), D_{\Delta}^2(\omega), R_1, R_2)$ space.

Following conventions often used to display results from oscillating gradient encoding the effects of restricted diffusion were quantified by a finite difference approximation of the rate of change of the diffusivity metrics with frequency within the investigated window, which in our case was $\omega_{\mathrm{max}} = 21$ Hz and $\omega_{\mathrm{min}} = 6.6$ Hz, and define

$$\Delta_{\omega/2\pi}E[D_{\mathrm{iso}}] = \frac{E[D_{\mathrm{iso}}(\omega_{\mathrm{max}})] - E[D_{\mathrm{iso}}(\omega_{\mathrm{min}})]}{(\omega_{\mathrm{max}} - \omega_{\mathrm{min}})/2\pi}, \tag{4}$$

and

$$\Delta_{\omega/2\pi}E[D_{\Delta}^2] = \frac{E[D_{\Delta}^2(\omega_{\mathrm{max}})] - E[D_{\Delta}^2(\omega_{\mathrm{min}})]}{(\omega_{\mathrm{max}} - \omega_{\mathrm{min}})/2\pi}. \tag{5}$$

**Diffusion-relaxation subdomains:** The $(D_{\mathrm{iso}}, D_{\Delta}^2, R_2)$ space was discretized into diffusion-relaxation subdomains defined by combinations of low, intermediate, and high regimes along each parameter dimension. Isotropic diffusivity was partitioned into three regimes corresponding to small, intermediate, and large microscopic length scales ($0.05 < D_{\mathrm{iso}} < 1.5$, $1.5 < D_{\mathrm{iso}} < 3$, and $D_{\mathrm{iso}} > 3$ $\mu$m$^2$/ms, respectively). Squared microscopic diffusion anisotropy was likewise partitioned into low, intermediate, and high regimes ($0 < D_{\Delta}^2 < 0.01$, $0.01 < D_{\Delta}^2 < 0.25$, and $D_{\Delta}^2 > 0.25$), reflecting increasing degrees of microscopic directional constraint. Transverse relaxation was partitioned into two regimes, corresponding to slow-relaxing ($1 < R_2 < 20$ s$^{-1}$) and fast-relaxing ($R_2 > 20$ s$^{-1}$) signal components, which are sensitive to differences in macromolecular content and water-tissue interactions. For each voxel, the following signal fractions were computed by summing the weights of consolidated components within the corresponding diffusion-relaxation subdomains, which we term here "microstructural phenotypes":

1. $|^{\ell}D_{\mathrm{iso}}|^{\hbar}D_{\Delta}^2|^{\ell}R_2|$: $0.05 < D_{\mathrm{iso}} < 1.5$ $\mu$m$^2$/ms; $D_{\Delta}^2 > 0.25$; $1 < R_2 < 20$ s$^{-1}$.
2. $|^{\ell}D_{\mathrm{iso}}|^{m}D_{\Delta}^2|^{\ell}R_2|$: $0.05 < D_{\mathrm{iso}} < 1.5$ $\mu$m$^2$/ms; $0.01 < D_{\Delta}^2 < 0.25$; $1 < R_2 < 20$ s$^{-1}$.
3. $|^{\ell}D_{\mathrm{iso}}|^{\ell}D_{\Delta}^2|^{\ell}R_2|$: $0.05 < D_{\mathrm{iso}} < 1.5$ $\mu$m$^2$/ms; $0 < D_{\Delta}^2 < 0.01$; $1 < R_2 < 20$ s$^{-1}$.
4. $|^{m}D_{\mathrm{iso}}|^{\hbar}D_{\Delta}^2|^{\ell}R_2|$: $1.5 < D_{\mathrm{iso}} < 3$ $\mu$m$^2$/ms; $D_{\Delta}^2 > 0.25$; $1 < R_2 < 20$ s$^{-1}$.
5. $|^{m}D_{\mathrm{iso}}|^{m}D_{\Delta}^2|^{\ell}R_2|$: $1.5 < D_{\mathrm{iso}} < 3$ $\mu$m$^2$/ms; $0.01 < D_{\Delta}^2 < 0.25$; $1 < R_2 < 20$ s$^{-1}$.
6. $|^{m}D_{\mathrm{iso}}|^{\ell}D_{\Delta}^2|^{\ell}R_2|$: $1.5 < D_{\mathrm{iso}} < 3$ $\mu$m$^2$/ms; $0 < D_{\Delta}^2 < 0.01$; $1 < R_2 < 20$ s$^{-1}$.
7. $|^{\hbar}D_{\mathrm{iso}}|D_{\Delta}^2|^{\ell}R_2|$: $D_{\mathrm{iso}} > 3$ $\mu$m$^2$/ms; $0 < D_{\Delta}^2 < 1$; $1 < R_2 < 20$ s$^{-1}$.
8. $|^{\ell}D_{\mathrm{iso}}|^{\hbar}D_{\Delta}^2|^{\hbar}R_2|$: $0.05 < D_{\mathrm{iso}} < 1.5$ $\mu$m$^2$/ms; $D_{\Delta}^2 > 0.25$; $20 < R_2 < 30$ s$^{-1}$.
9. $|^{\ell}D_{\mathrm{iso}}|^{m}D_{\Delta}^2|^{\hbar}R_2|$: $0.05 < D_{\mathrm{iso}} < 1.5$ $\mu$m$^2$/ms; $0.01 < D_{\Delta}^2 < 0.25$; $20 < R_2 < 30$ s$^{-1}$.
10. $|^{\ell}D_{\mathrm{iso}}|^{\ell}D_{\Delta}^2|^{\hbar}R_2|$: $0.05 < D_{\mathrm{iso}} < 1.5$ $\mu$m$^2$/ms; $0 < D_{\Delta}^2 < 0.01$; $20 < R_2 < 30$ s$^{-1}$.

### Supplementary Figures

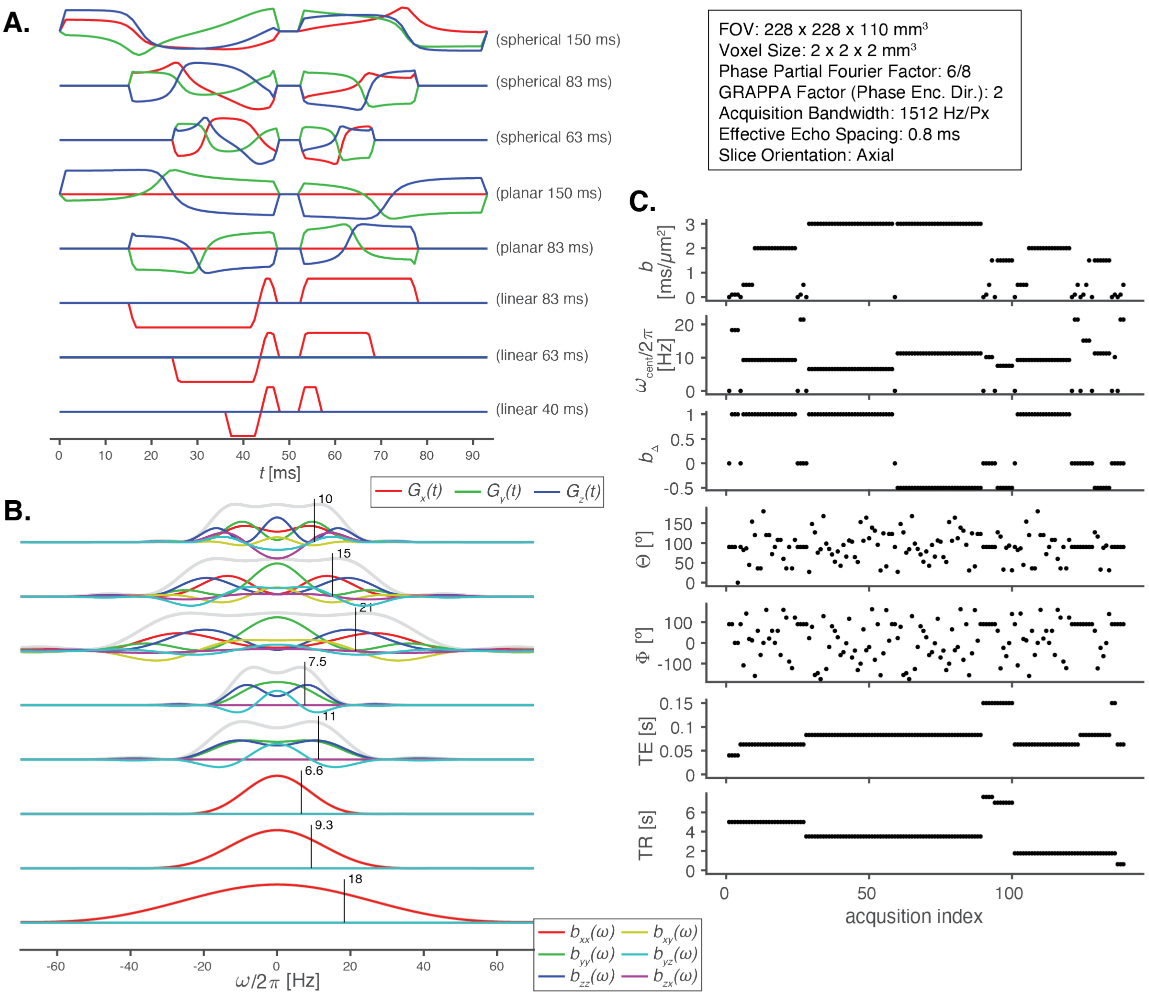


Figure S1: **MD-MRI acquisition and encoding design** (A) Time-dependent optimized diffusion gradient waveforms $G_x(t)$, $G_y(t)$, and $G_z(t)$ used to generate linear, planar, and spherical b-tensors at different echo times. (B) Corresponding diffusion spectra $b(\omega)$, illustrating the broad frequency content of the encoding waveforms. (C) Acquisition protocol showing the repetition time (TR), echo time (TE), b-tensor magnitude ($b$), normalized anisotropy ($b_\Delta$; planar: $-0.5$, spherical: 0, linear: 1), gradient orientation angles $(\theta, \phi)$, and centroid frequency $\omega_{cent}/2\pi$, plotted as a function of acquisition index.

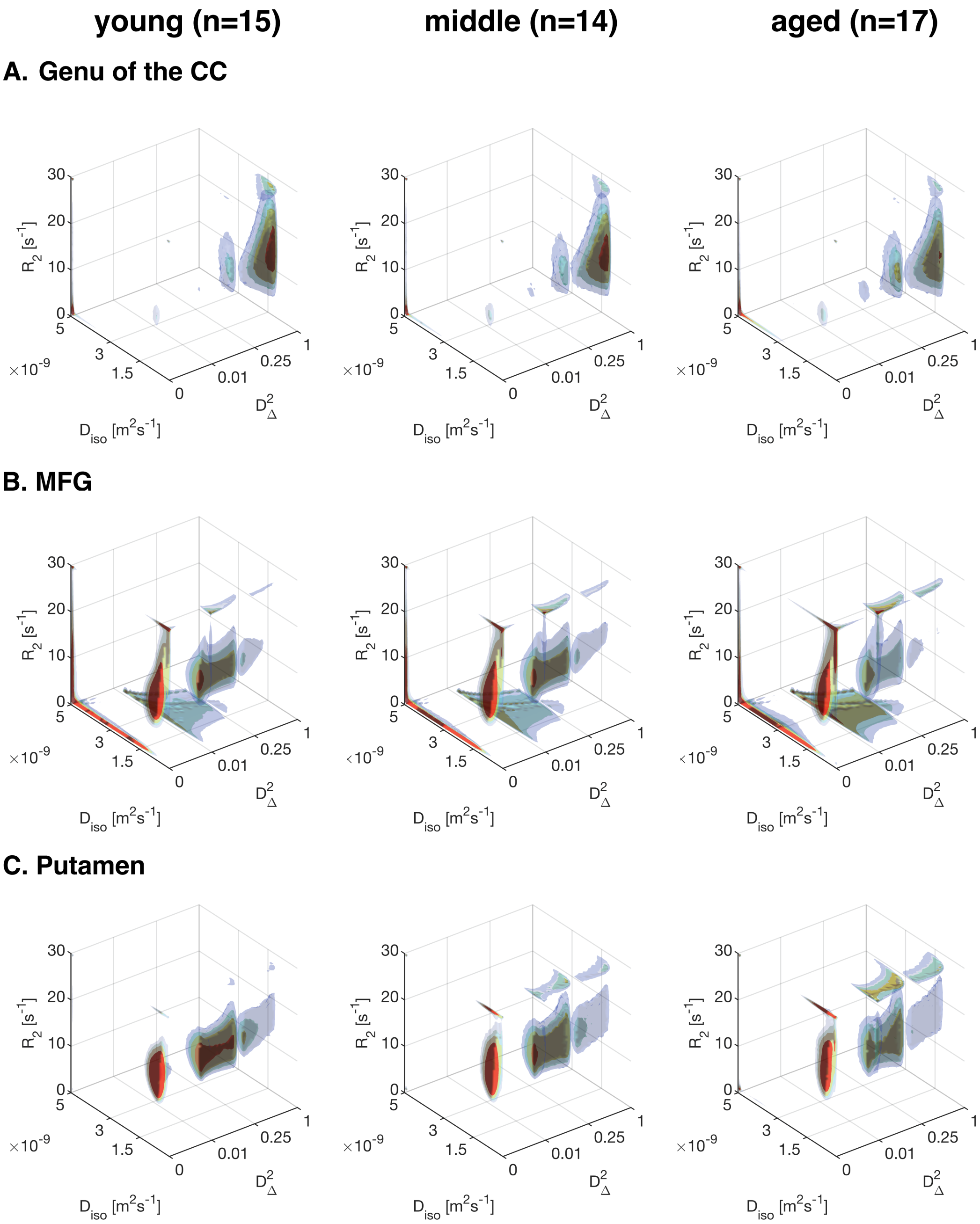


Figure S2: Age-related redistribution of joint diffusion-relaxation phenotypes across major brain tissue classes. Group-averaged three-dimensional joint $D_{\text{iso}}$–$D_{\Delta}^{2}$–$R_2$ distributions projected at maximal frequency, $\omega_{min}$ = 21 Hz, are shown for young (left), middle-aged (middle), and aged adults (right) in the (A) genu of the corpus callosum, (B) middle frontal gyrus (MFG), and (C) putamen. Across brain regions, aging is marked by multilateral microstructural shifts that reflect complex trajectories and together form a shared aging signature.

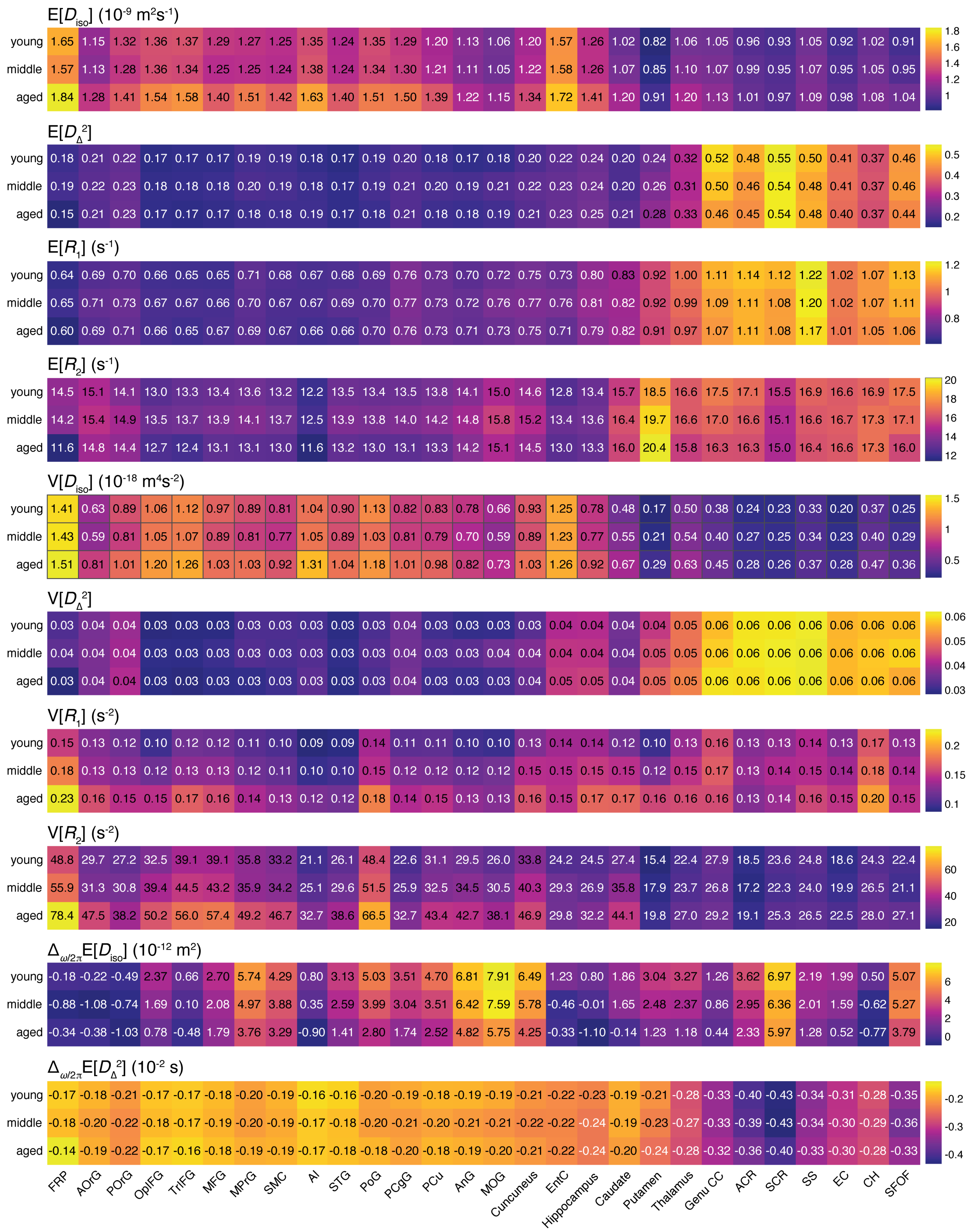


Figure S3: Region-of-interest–averaged MD-MRI diffusion–relaxation parameters, stratified by age group.

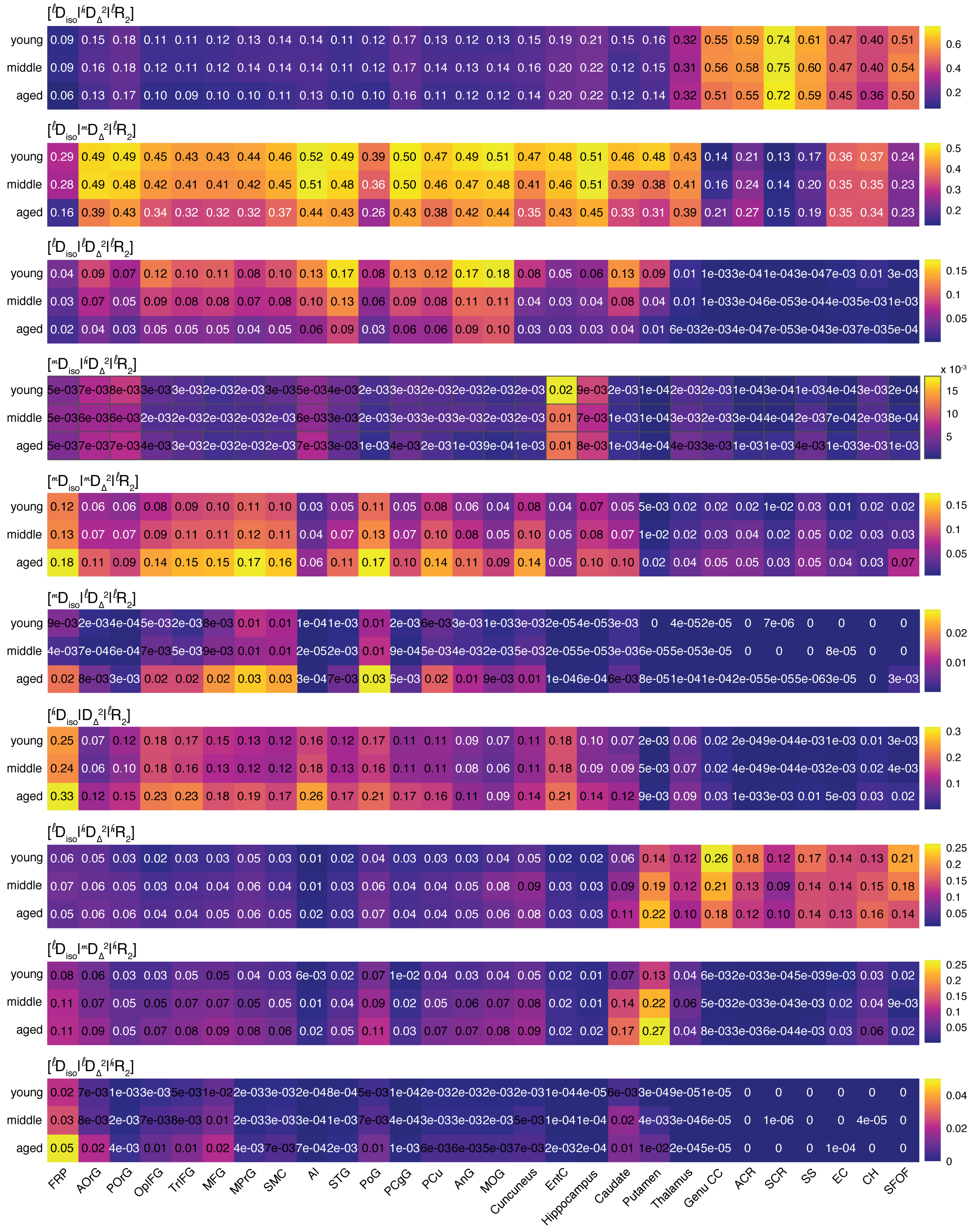


Figure S4: Region-of-interest–averaged MD-MRI microstructural phenotypes, stratified by age group. Some phenotypes have clear regional specificity; for example, both $|^{\ell}D_{\rm iso}|^{\hbar}D^2_{\Delta}|^{\ell}R_2|$ and $|^{\ell}D_{\rm iso}|^{\hbar}D^2_{\Delta}|^{\hbar}R_2|$ are more prevalent in WM. Conversely, lower anisotropy subdomains, e.g., $|^{\ell}D_{\rm iso}|^{\ell}D^2_{\Delta}|^{\ell}R_2|$ or $|^{m}D_{\rm iso}|^{m}D^2_{\Delta}|^{\ell}R_2|$, mainly dominate cortical and subcortical regions.

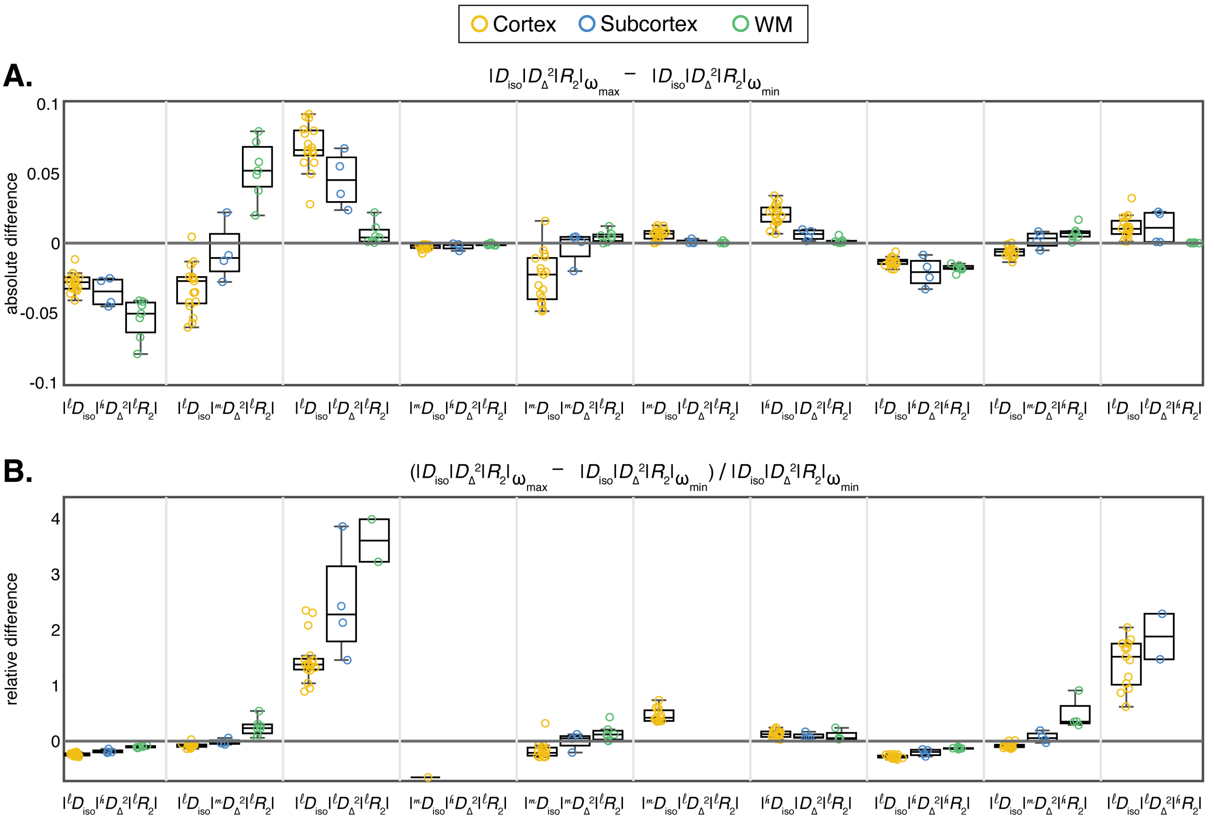


Figure S5: Diffusion frequency dependence of microstructural phenotypes. Panels A and B show the absolute and relative differences, respectively, between high- and low-frequency estimates averaged across the aged adult group. Differences are grouped and color-coded by brain region (cortical GM, subcortical GM, and WM). Phenotypes with ROI-averaged intensity under 0.01 were omitted from the plot to avoid bias.

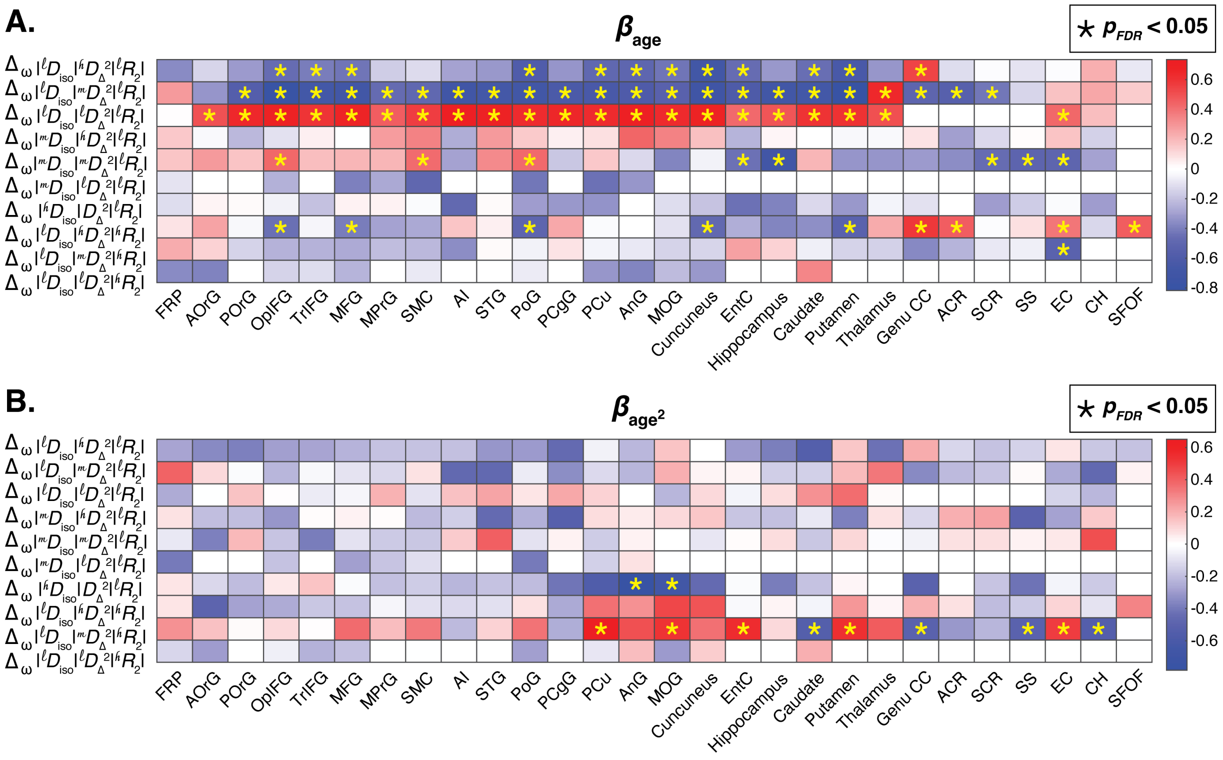


Figure S6: Linear and nonlinear age associations of frequency dependence of microstructural phenotypes across brain regions. (A) Standardized linear age coefficients ($\beta_{\text{age}}$) estimated for signal fraction frequency dependence associated with distinct diffusion-relaxation subdomains across cortical, deep GM, and WM ROIs. (B) Corresponding quadratic age coefficients ($\beta_{\text{age}^2}$) capturing nonlinear lifespan trajectories. Phenotypes with ROI-averaged intensity under 0.01 were omitted from the plot to avoid bias.

[11] Markus Nilsson, Filip Szczepankiewicz, Björn Lampinen, André Ahlgren, Jo ao P. de Almeida Martins, Samo Lasic, Carl-Fredrik Westin, and Daniel Topgaard. An open-source framework for analysis of multidimensional diffusion mri data implemented in matlab. Proceedings of the International Society for Magnetic Resonance in Medicine, Paris, France, 2018.
